# Early female germline development in *Xenopus laevis*: stem cells, nurse cells and germline cysts

**DOI:** 10.1101/2025.07.29.667541

**Authors:** Asya Davidian, Allan C Spradling

## Abstract

In Drosophila, germline cysts arise through synchronous mitotic divisions and acquire a polarized architecture organized by the fusome, which guides oocyte specification and supports meiotic progression. Similar cyst structures exist in non-mammalian vertebrate ovaries, but their polarity and function have remained uncertain. Using single-cell RNA sequencing and high-resolution imaging, we reconstructed the germ cell differentiation trajectory in *Xenopus laevis* and uncovered striking parallels with invertebrate and mouse cyst development. We identified a distinct germline stem cell (GSC) population marked by piwil4, low translational activity, and expression of neuronal-specific and transposon-silencing genes. Downstream from GSCs, during cyst development, an asymmetric fusome-like structure (FLS) composed of stable microtubules forms a rosette-like connection between cystocytes and co-localizes with Golgi vesicles and ER, suggesting polarized trafficking. In contrast to previous claims, ∼90% of EdU-prelabeled cyst cells turned over rather than forming oocytes, consistent with a nurse cell fate. The striking parallels described here between cyst and fusome formation, polarization, cyst breakdown, and nurse cell turn over to produce relatively few oocytes, argue that amphibian cysts have important functions in female gametogenesis.

## INTRODUCTION

A stereotyped program of synchronous mitotic cell cycles with incomplete cytokinesis that produces germline cysts arose at the dawn of animal evolution and remains a feature of male gametes in all animal groups. Female gametogenesis also utilizes germline cysts in humans, other mammals as well as many other vertebrate and invertebrate groups (reviews: Lu et al., 2017; Chaigne and Brunet, 2022; Gerhold et al., 2022; Spradling, 2024; Brubacher, 2024). In Drosophila, cysts arise from germline stem cells (GSCs) located at the anterior tip of the germarium. GSCs divide asymmetrically to produce a cystoblast, which then undergoes four synchronized divisions to generate a 16-cell cyst linked by a cytoskeleton-rich structure known as the fusome, that passes through the ring canals joining the cells. Cysts remain during early meiotic prophase in females, where they give rise in addition to oocytes, to a second cell type, nurse cells, that are known or proposed to contribute to oocyte development. In Drosophila and other higher insects, cysts have been recognized for more than 100 years, and genetic studies show Drosophila cysts are essential for oocyte production and fertility.

The biology of the Drosophila fusome has been extensively analyzed (reviewed in King, 1970; Hinnant et al. 2020; Spradling et al. 2022) and compared to other insects (Büning, 1994). The fusome arises from mitotic spindle remnants during cyst formation and segregates asymmetrically at each division reflecting the developing polarity of the cyst. After meiotic entry, the fusome reaches its maximum diameter at the start of meiosis and changes morphologically and in protein content throughout meiotic prophase (Grieder et al. 2000; Lighthouse et al., 2008). The fusome turns over at the time of follicle formation, after ushering the movement of organelles and other materials from the nurse cells into the oocyte and its forming Balbiani body (Bb) (Cox and Spradling, 2003; Cox and Spradling, 2006).

Three types of cyst functions have been identified. Nurse cells acquired their name from their function in higher insects, where they synthesize and transfer nearly all oocyte materials and assist oocytes in some species to grow exceptionally large. Cysts in a wide range of species in both males and females also serve a defensive function by sharing small RNAs targeting transposable elements and other parasites attempting to expand in germ cells (Lim et al. 2014; Chen and Aravin, 2023; Sorkin et al., 2025). Finally, germ cell rejuvenation processes needed for species perpetuation take place during meiosis in *Saccharomyces cerevisiae* (review: Goodman et al. 2020) and probably all eukaryotes. Metazoans likely carry out major aspects of rejuvenation during meiosis in female gametogenesis. Drosophila germline cysts facilitate the rejuvenation of mitochondria, and potentially other organelles that are collected in the Balbiani body (Cox and Spradling, 2003; Cox and Spradling, 2006; Marlow and Mullins, 2008; Lieber et al. 2019; Palozzi et al. 2022; Spradling et al. 2022; Yamashita, 2023; Monteiro et al. 2023).

Studies of Xenopus have contributed immensely to our understanding of oogenesis and its role in embryonic development. Xenopus oogenesis has been well described and used to study rDNA amplification and many other aspects of oocyte development and lampbrush chromosome function (Gall, 1968; Gall and Pardue, 1969; Al-Mukhtar and Webb, 1971; Coggins and Gall, 1972; Dumont, 1972). Xenopus oocytes form a prominent Balbiani body that initially arises at stage 1 and like the zebrafish Bb persists for much of oogenesis, unlike the transient Bbs of mice and Drosophila. The frog Bb plays a role in localizing mRNAs initially at the vegetal pole that contribute to embryo patterning and to germ plasm formation (Kloc et al. 2004b; Aguero et al. 2017). Studies of germline cysts in Xenopus ovaries were pioneered by Kloc et al. (2004a) who described 16-cell cysts, formed in synchronous mitotic divisions, containing typical ring canals as visualized by electron microscopy. Serial section reconstruction of developing cysts showed possible evidence of a fusome-like structure. However, Xenopus cysts were not observed to undergo apoptosis leading the authors to speculate that all cyst cells develop into oocytes.

The existence of GSCs in the Xenopus ovary remains unresolved. Although a unique cell type with a large nucleus and mitochondrial cloud—termed the “primary oogonial cell”—was described decades ago (Al-Mukhtar and Webb, 1971; Tourte, 1981), the prevailing view is that amphibians, like mammals, produce a finite pool of oocytes early in life (Callen et al., 1986; Ogielska et al., 2013; Ogielska et al., 2024). Small oogonial patches in adult Rana ovaries have been reported but are considered non-functional (Ogielska et al., 2013). In contrast, adult GSCs sustaining oogenesis have been well documented in zebrafish and medaka (Nakamura et al., 2010; Beer and Draper, 2013; Liu et al., 2022), and recent scRNA-seq studies have enabled their molecular identification (Liu et al., 2022). These findings highlight the need to re-examine the Xenopus ovary using modern tools to assess whether a bona fide GSC population exists.

Germline cysts have been identified in mice (Pepling and Spradling, 1998), Xenopus (Kloc et al 2004a), fish (Marlow and Mullins, 2008) and birds (Skalko et al., 1972; Dong et al., 2022), and in mice they produce both oocytes and nurse cells (Lei and Spradling, 2016; Niu and Spradling, 2022). Nonetheless, the idea that non-mammalian vertebrates generate polarized germline cysts to specify oocytes and nurse cells, and to support oocyte development has remained controversial (review: Spradling et al. 2022). Rather, the idea that vertebrate cysts are present but have no conserved function continues to be maintained especially in the case of lower vertebrates such as amphibians and fish (see Brubacher, 2024).

In this study, we examined the development of female germ cells in *Xenopus laevis* to look for GSCs and to better characterize the biology of amphibian germline cysts. Using single-cell transcriptomics and high-resolution microscopy we identified a distinct population of GSCs. These cells expressed a distinctive program of transcription marked by *piwil4*, neuronal and transposon-silencing gene signatures. We show that germline cysts assemble a fusome-like structure (FLS) composed of stable microtubules which are often associated with Golgi vesicles and ER. Microtubules were unevenly distributed within cysts, suggesting the emergence of early structural asymmetry. Furthermore, we demonstrate that the vast majority of cyst cells do not become oocytes, but instead exhibit characteristics of nurse cells, persisting transiently before turnover and displaying low UMI counts, similar to nurse cells previously described in mice (Niu and Spradling, 2022). However, unlike fly and mouse nurse cells, evidence of organelle bulk transfer was not confirmed. These findings support a largely conserved role for germline cysts and nurse cells in Xenopus, and suggest that early cyst polarity, possible selective oocyte formation, and nurse-like cell turnover are ancestral features of female gametogenesis across animals.

## RESULTS

### Germline differentiation trajectory revealed by scRNA-seq and cytological markers

To investigate early female germline development in *Xenopus laevis*, we performed single-cell RNA sequencing (scRNA-seq) on post-metamorphic ovaries which are enriched for germ cells undergoing cyst formation and meiotic entry. Transcriptomes were generated from 18,410 cells, including 8,544 germline and 9,866 somatic cells, and processed using Seurat. Germline cells were identified based on expression of conserved markers such as *ddx4* and *dazl*, and subsequently reclustered to resolve transcriptional heterogeneity. UMAP analysis revealed a continuous developmental trajectory beginning with a distinct germline stem cell (GSC) cluster and progressing through cyst formation, meiotic prophase, and early follicle stages (Figure 1A). To annotate these clusters, we examined the expression of stage-specific markers (Figure 1B, C) and integrated morphological features from EdU and telomere DNA FISH experiments (Figure 1D).

**Figure 1.**
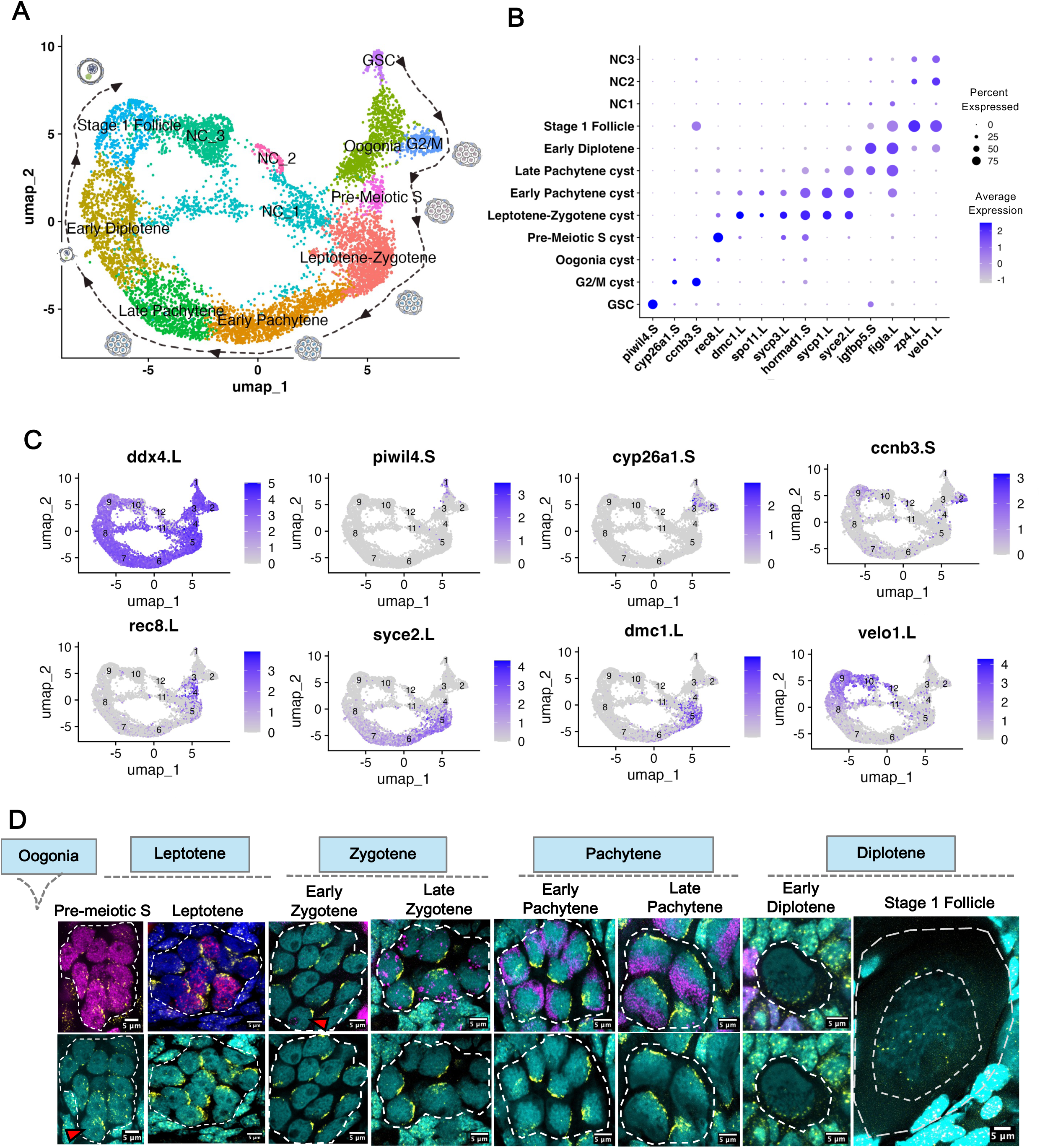
Development of cysts based on scRNAseq and Telomere FISH. **A-** UMAP visualization of 11 transcriptionally distinct germline clusters identified from integrated scRNA-seq data of juvenile Xenopus ovaries across three biological replicates. Developmental progression follows a clockwise trajectory along the outer ring, starting from germline stem cells (GSCs) through proliferating oogonia, premeiotic S-phase, meiotic cyst stages (leptotene, zygotene, pachytene), and culminating in early diplotene primordial follicles and stage I follicles. The inner ring comprises transcriptionally distinct nurse-like cell populations (NC1) corresponding to different meiotic cyst stages, and additional clusters (NC2 and NC3) likely representing “wave 1” follicles. **B** – Dot plot comparing expression patterns of selected marker genes across different germ cell clusters. **C** – Feature plots showing marker gene expression mapped onto UMAP coordinates. **D** – Developmental progression of Xenopus germline cysts visualized by EdU labeling (1.5 h, magenta), telomere FISH (yellow) and DAPI (cyan). In the pre-meiotic S-phase, telomeres cluster at the nuclear envelope, with early bouquet formation in “leading” cystocytes (arrowhead). Leptotene/zygotene stages show bouquet formation and the onset of rDNA amplification (arrowhead). During pachytene, nuclear size increases and rDNA amplification continues, forming an “rDNA cap.” In early diplotene, cyst fragments form primordial follicles, and telomeres begin to disperse. Stage I follicles display large germinal vesicles (GV) with forming lampbrush chromosomes and telomere detachment.

Just downstream of the GSC cluster, cells expressed *cyp26a1* (cytochrome P450 family 26 subfamily A member 1; Feng et al., 2015), a meiotic entry repressor, consistent with oogonia forming germline cysts. A neighboring G2/M cluster, characterized by high expression of *ccnb3* (cyclin B3), *cdk1* (cyclin-dependent kinase 1), *melk* (maternal embryonic leucine zipper kinase) and *mki67* (marker of proliferation ki-67), represented actively dividing cells (Badouel et al., 2010; Lara-Gonzalez et al., 2024). The pre-meiotic S cluster, marked by *rec8* (meiotic recombination protein; Watanabe et al., 2001), corresponded to cystocytes undergoing DNA replication, confirmed by EdU labeling and the onset of telomere clustering. Cells expressing *spo11*, *dmc1*, *sycp3*, and *hormad1*—key genes involved in meiotic double-strand break formation, recombination, and synaptonemal complex assembly—were annotated as leptotene–zygotene cysts (Carofiglio et al., 2018). These cells exhibited prominent telomere bouquets and showed early nuclear polarization, characterized by the repositioning of the nucleolus and the initiation of rDNA amplification. During early pachytene, elevated expression of the same markers and *syce2* (synaptonemal complex central element protein 2) coincided with the formation of a prominent rDNA cap opposite the telomeres. The late pachytene cluster, defined by continued expression of *syce2* and *hormad1*, aligned with increasing nuclear size and a fully developed rDNA cap. In early diplotene, rDNA amplification ceased, telomeres began to disperse, and *velo1* was expressed—a marker of the Balbiani body (Claussen and Pieler, 2004; Divyanshi and Yang, 2023). Induction of *figla*, an oocyte-specific transcription factor (Liang et al., 1997), marked the transition to pre-follicular oocytes. Finally, cells expressing *zp4* (zona pellucida; Bleil and Wassarman, 1980) and *velo1* were annotated as stage 1 follicles (Dumont, 1972) with enlarged germinal vesicles and telomeres detached from the nuclear membrane.

On UMAP we observe an outer ring of canonical germline progression in clockwise direction, along with an inner ring which we named NC1, NC2 and NC3. The cells in the NC clusters show low UMI values (Figure 7D) compared to clusters on outer ring, similar to mouse nurse cells with low UMI due to the transfer of cytoplasm to pro-oocytes (Niu and Spradling, 2022). Xenopus nurse cells in cluster NC1 arise early in meiosis and showed many similarities to early mouse nurse cells, for example in reduced expression of the key RNA regulator *daxl* (0.25 of leptotene/zygotene) (Suppl. Figure S1A). Other downregulated NC1 genes (also affected in E14.5 mouse NC cluster 19, Niu and Spradling, 2022) were a group of meiosis-specific genes: *sycp1, sycp3, syce2, dmc1, spata22* (all 0.1-0.3), cilia regulator *arl3* (0.22), *hormad1* (0.15), and the chromatin regulator *macroh2a2* (0.212). Overexpressed genes in NC1 include *42sp43* and *42sp50*, a Xenopus oocyte specific translational regulatory complex (8-12X), *tmsb15* (7.6) and *tmsb4x (*4.0) both thymosin beta proteins implicated in sequestering g-actin, and other genes with potential effects on nurse cell function. In contrast to NC1 downregulated genes, the expression in mouse oogenesis of NC1 over-expressed were mostly not detected.

Multiple genes down and up-regulated in NC2 and NC3 were also examined relative to expression in stage 1 follicles. A sample of upregulated genes included *ddx25* (5.1, 5.6, i.e. NC2, NC3), DEAD box helicase involved in RNA based transcript regulation in the testis, whose function in the ovary is little studied, *rab11fip1* a rab11endosome trafficking modulator (1.7, 5.3), *larp6* (2.2, 4.1) a translational regulator, *kif20b* (3.0, 3.6) a microtubule motor and meiotic spindle regulator. Downregulated genes include *acp5* (acid phosphatase (0.016, 0.39)) whose ovarian function is unknown, *not* (notochord homeobox) (0.097, 0.65) and others. These findings show that the expression of genes in nurse cells is highly regulated, and in some cases changes are conserved from mouse to Xenopus. Our hypothesis is that NC2 and NC3 germ cells derive from oocytes of a “wave 1” subpopulation of Xenopus follicles that are programmed to turn over, while their somatic cells carry out a specialized function (Yin and Spradling, 2025).

### Identification and properties of a Xenopus female GSC population

At the origin of the germline trajectory, we identified a transcriptionally distinct cluster defined by high and specific expression of *piwil4*. Using hybridization chain reaction fluorescence *in situ* hybridization (HCR-FISH) with *piwil4.S* RNA probe (Figure 2A-A’’), we validated these cells as large, *ddx4*-positive (Figure 2C-C’), individual germ cells with multilobed nuclei preferentially localized near the ovarian surface. This cell type closely resembles the “primary oogonial cells” previously described morphologically in Xenopus (Al-Mukhtar and Webb, 1971). We also detected similar *piwil4*-expressing cells in the adult ovary, specifically within the ovarian epithelium (Suppl. Figure S2). In both juvenile and adult ovaries, these cells exhibit characteristic features of undifferentiated germ cells: their nuclei are large and polymorphic, with relatively open chromatin as shown by DAPI staining (Figure 2A”), they display a prominent mitochondrial cloud (Figure 2B, E) localized around the centrosome and an enriched endoplasmic reticulum (ER) cluster (Figure 2B, D), reminiscent of the spectrosome found in Drosophila GSC.

**Figure 2.**
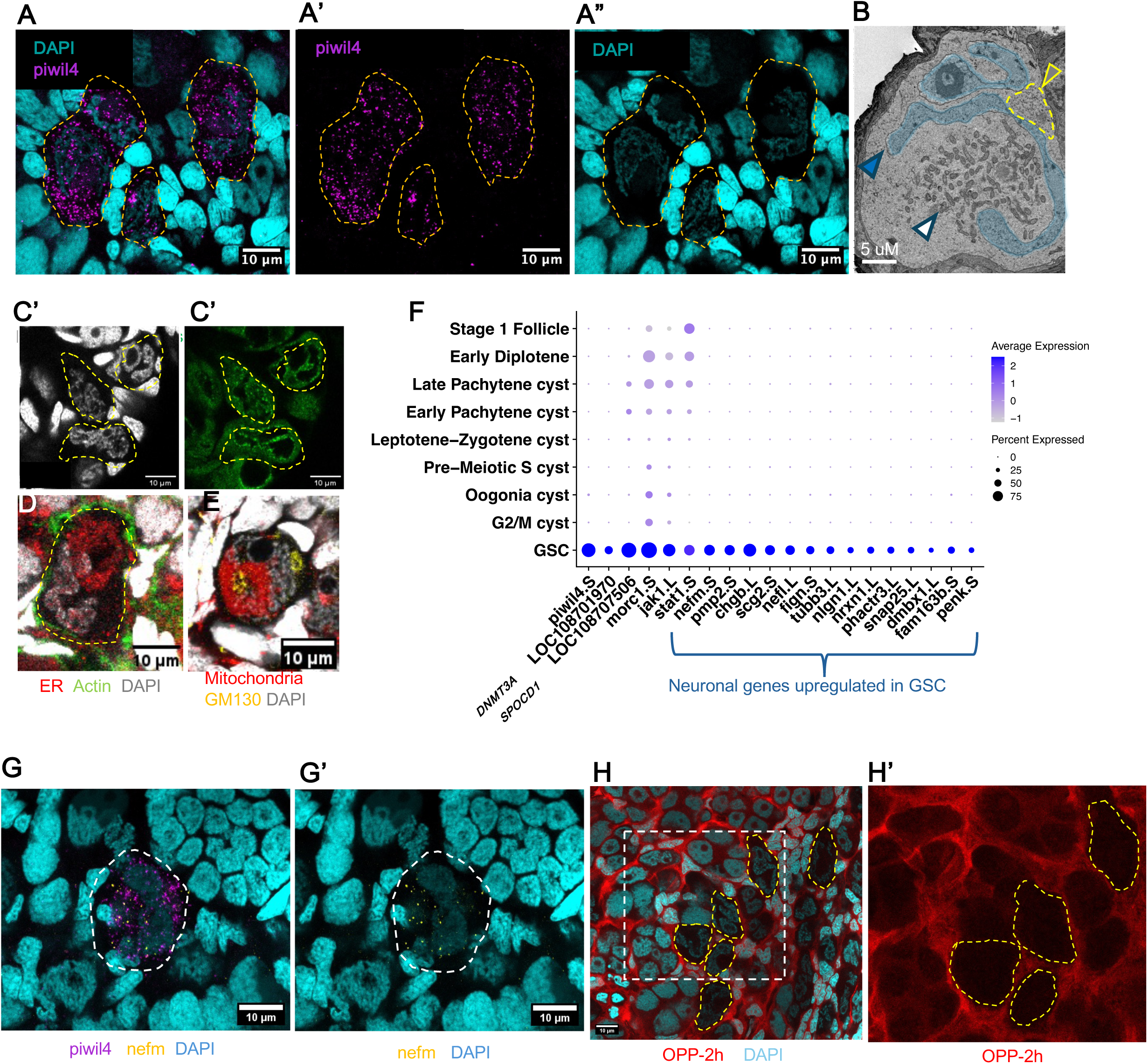
Identification of GSCs in Xenopus ovary. **A** – Hybridization chain reaction (HCR) RNA-FISH for *piwil4.S* (magenta) combined with DAPI staining (cyan) in whole-mount juvenile Xenopus ovary. **B** – GSC under electron microscopy showing a polymorphic nucleus (blue highlight), mitochondrial cloud (white arrowhead), and ER-like structures (yellow arrowhead). **C-C’** – GSCs stained with Ddx4 (green) and DAPI (gray). **D** – GSC stained with PDI (red), phalloidin (green), and DAPI (gray). **E** – GSC stained for mitochondria (citrate synthase, red), Golgi (GM130, yellow), and DAPI (gray). **F** – Dot plot comparing expression patterns of selected marker genes for GSCs across different germ cell clusters. **G-G’** – Double HCR RNA FISH for *piwil4.S* (magenta) and *nefm.S* (yellow) combined with DAPI (cyan) in whole-mount juvenile Xenopus ovary. **H-H’** – Whole-mount juvenile Xenopus ovary stained with OPP (2 hr, red) and counterstained with DAPI (cyan). **H’** shows a magnified view of the dashed rectangle in panel H. Quiescent GSCs are outlined with yellow dashed lines.

Molecularly, this cluster also expressed *spocd1* (LOC108707506) and *morc1* (Figure 2F), two core effectors of the PIWI–piRNA pathway involved in transposon silencing, previously shown to be critical in mouse gonocytes (Uneme et al., 2024). Their co-expression with *piwil4* suggests that Xenopus GSCs initiate transposon repression early in germline development. Additional markers, including *dnmt3a* (LOC108701970, DNA methyltransferase 3a) and *chgb* (chromogranin B) – also reported in zebrafish GSCs (Liu et al., 2022) – were highly enriched in this cluster. We also observed upregulation of the JAK1–STAT1 signaling pathway – known for its conserved role in stem cell regulation across species (Stine and Matunis, 2013) – enriched in this cluster (Figure 2F).

Unexpectedly, the cluster showed strong enrichment for genes typically associated with neuronal function (Figure 2F, Suppl. Figure S3). These included cytoskeletal and synaptic components (*neurofilament medium and light chains, βIII-tubulin, neuroligin 1, neurexin 1*), neuropeptide precursors and secretory factors (*VGF, secretogranin II, proenkephalin, chromogranin, synuclein beta, neuropeptide y, synaptosome-associated protein*), and regulators of neuronal identity (*fam163, diencephalon/mesencephalon homeobox)* (Figure 2F, Suppl. Figure S3). Co-expression of *piwil4* and *nefm* was confirmed by double HCR-FISH (Figure 2G-G’).

In line with their stem-like status, GSCs also appeared to be translationally quiescent compared to downstream germ cells. This was evident from low O-propargyl-puromycin (OPP) incorporation (Figure 2H-H’), indicating reduced global protein synthesis, and was supported by scRNA-seq data, where the downstream oogonia clusters showed marked upregulation of different ribosomal protein genes (Suppl. Figure S3). Such translational repression is a hallmark of many stem cell populations and has been implicated in preserving genome integrity and stemness (Saba et al., 2021).

To test whether adult Xenopus can utilize GSCs to replenish the oocyte pool, we examined adult females several years after partial ovariectomy. We observed small, regenerated ovarian lobes (Suppl. Figure S1 C-C”) that contained germline cysts at various stages—including dividing oogonia, meiotic cysts, and stage I–II follicles—as well as cells with features consistent with GSCs. In contrast, such structures were not found in frogs that had not undergone ovariectomy. These findings suggest that the adult Xenopus ovary retains regenerative capacity and harbors a GSC population capable of reinitiating oogenesis following tissue loss.

### Developing cysts are stably interconnected by a microtubule-rich fusome-like structure

Downstream of the GSC population, germline cysts begin to form through mitotic divisions of oogonia and early meiotic entry, primarily along the ovarian periphery (Coggins, Gall 1972; Kloc et al, 2004). To identify and study developing cysts, ovaries from 50-60 day-old froglets were labeled with EdU and analyzed as whole mounts. Using 3D reconstruction in Imaris, we identified clusters of 2-, 4-, 8-, 16-, and 32 EdU-positive cells as cysts (Suppl. Figure S4A-B). However, as many as ∼17% of cysts did not contain 2ⁿ cells, such as 6-, 19-or 25 cells (Suppl. Figure S4A-B), suggesting that cyst fragmentation occurs, similar to observations in mice (Lei and Spradling, 2013; Levy et al. 2024). An example of such a cyst fortuitously labeled with EdU and ring canals (via Kif23, a kinesin family member and component of the centralspindlin complex) as an 8-cell cyst (Figure 3A) now shows two closely associated but separate 6-cell and 2-cell groups with no ring canal between them. Live imaging of dissected ovaries stained with Hoechst revealed that cystocytes exhibit active crawling-like movement which may contribute to cyst breakage (Figure 3B; Suppl. Figure S4C). This type of germline cell behavior was observed especially during oogonia stages before meiotic prophase. In addition, unlike the perfect synchrony observed in Drosophila germ cell cysts, Xenopus cysts showed limited but detectable asymmetry in cell cycle progression with distinguishable “leader” and follower cells as observed in both fixed and live samples (Suppl. Figure S4D, E).

**Figure 3.**
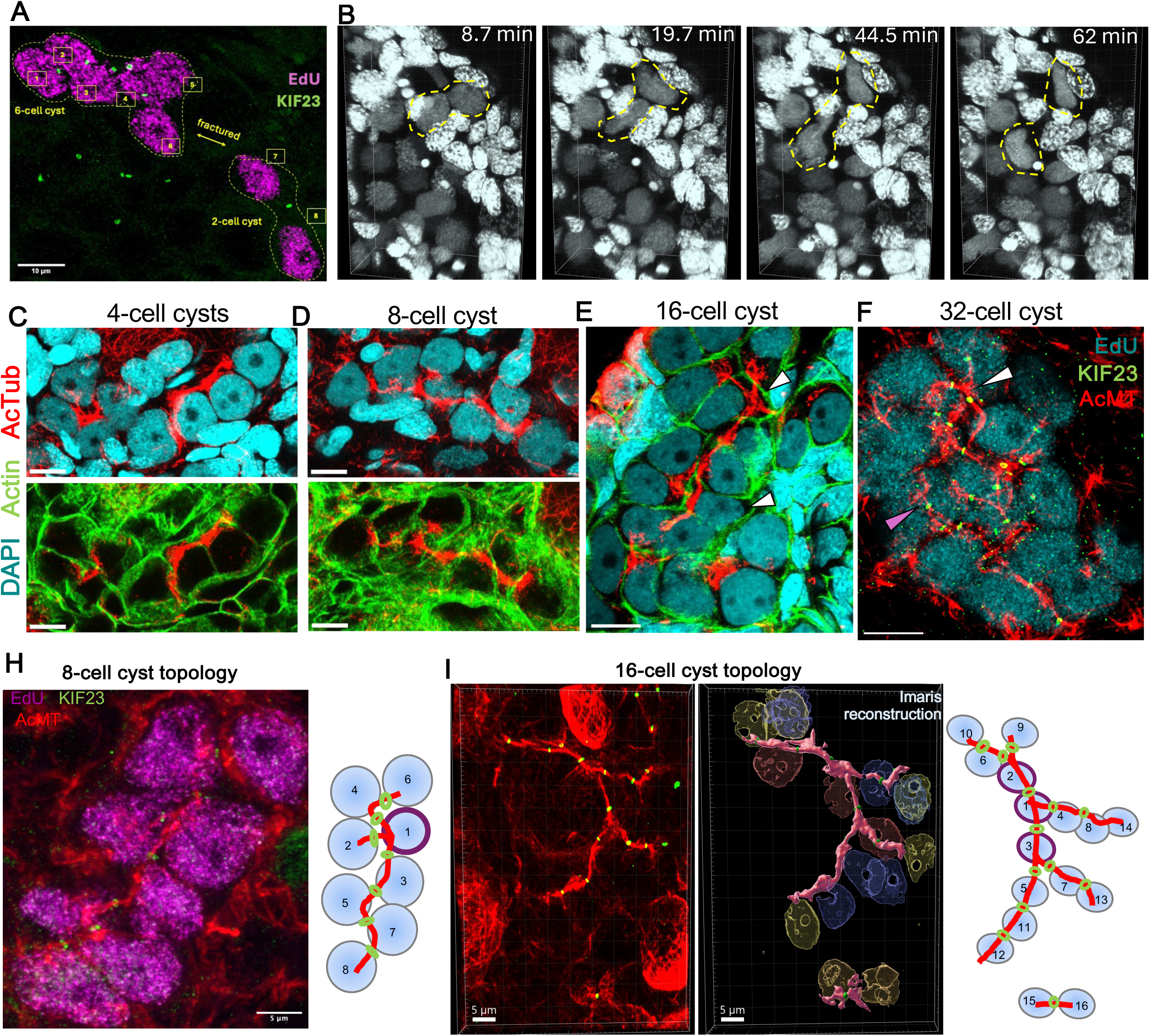
Germline cysts and their microtubule-based fusome like structure. **A** – Germline cyst labeled with EdU (∼2 hr, magenta) and Kif23 antibody (ring canals, green) undergoing fragmentation into a 6-cell group and 2 separate cells. **B** – Live imaging (∼2 hr) of a juvenile ovary stained with Hoechst (gray), showing motile cystocytes (yellow dashed lines). **C–E** – 4-, 8-, and 16-cell cysts stained with acetylated microtubules (red), phalloidin (green), and DAPI (cyan). **F** – 32-cell cyst labeled with EdU (cyan), acetylated microtubules (red), and Kif23 (green). **H** – Topology of an 8-cell cyst labeled with EdU (magenta), Kif23 (green), and acetylated microtubules (red), with adjacent cartoon illustrating the intercellular connectivity. **I** – Fragmented 16-cell cyst (14-cell group + 2 cells) labeled with Kif23 (green) and acetylated microtubules (red); adjacent Imaris 3D reconstruction showing cyst nuclei and FLS (pink), alongside a cartoon of the actual interconnections.

A major feature of Drosophila cyst is the fusome - an asymmetric, microtubule-rich organelle containing smooth ER cisternae and more than 20 identified proteins (review: Spradling et al. 2022). A strikingly similar structure was observed in Xenopus cysts stained for stable microtubules (Figure 3C-F). Immunostaining of ovaries for acetylated tubulin, actin, and DAPI revealed that bundles of stable microtubules (MT) traverse the sister cells in a branched path within 8- and 16-cell cysts (Figure 3C-E). These MT bundles pass through intercellular bridges (ring canals) as shown in a 32-cell cyst labeled for Kif23, a kinesin-like protein 23 (Figure 3F).

Interestingly, microtubules are unevenly distributed within the cyst: cells with multiple intercellular connections often exhibit thicker microtubule bundles, suggesting a polarized organization of the cyst (Figure 3E, F; white arrowheads indicate cells with 3 or 4 ring canals and prominent microtubule bundles; pink arrowhead marks a cell with thinner bundles). Due to their resemblance to the Drosophila fusome, we refer to these structures as the “fusome like structure (FLS).” Like the fusome, FLS exhibits branching, although Xenopus cysts show less branching than mazimum branching feature of Drosophila cysts. Due to the compact architecture of Xenopus cysts full topological reconstruction is rarely possible. Nonetheless, in well-preserved examples, we were able to analyze cyst organization and identify key differences from Drosophila. In 8-cell cysts, only one cell usually had three ring canals, compared to two such cells in fly cysts (Figure 3H). In a fragmented 16-cell cyst, we observed a split into 14- and 2-cell groups, and within the 14-cell portion, three cells each had three ring canals—unlike the Drosophila pattern of two cells with four connections (Figure 3I). These findings suggest that while Xenopus cysts share overall architecture with insect cysts, their topology and fragmentation dynamics are distinct. Notably, the FLS did not stain with spectrin, actin, or the Drosophila hts antibody (1B1) (data not shown), indicating possible molecular divergence despite morphological similarity.

### Germline cysts and their FLS during mitotic and early meiotic stages

The fusome in well studied systems arises from the metaphase spindles following cystocyte mitotic division, whose turnover is arrested by the failure to complete cytokinesis (Mathieu et al. 2022; Price et al. 2023; Glotzer, 2025). In forming Xenopus cysts, spindles appeared normal at metaphase (Figure 4A). In late telophase, spindle microtubules contract and bundle within the intercellular bridge, forming the midbody (Figure 4B). Subsequently, these midbodies within the cyst become stabilized, giving rise to stable intercellular bridges with no microtubule gap within bridges (Figure 4C). Exactly how this stabilized midbody leads to a permanent intercellular bridge and altered microtubule polarity is not well understood. As the cyst enters interphase, microtubule bundles interconnect all cystocytes, forming a network in which stable microtubules contribute to cystocyte connectivity (Figure 4D). While all EdU-positive cysts (S-phase) show strong microtubule interconnections (Suppl. Figure S5), many EdU-negative cysts with the same morphology (interphase) also maintain robust microtubule interconnections.

**Figure 4.**
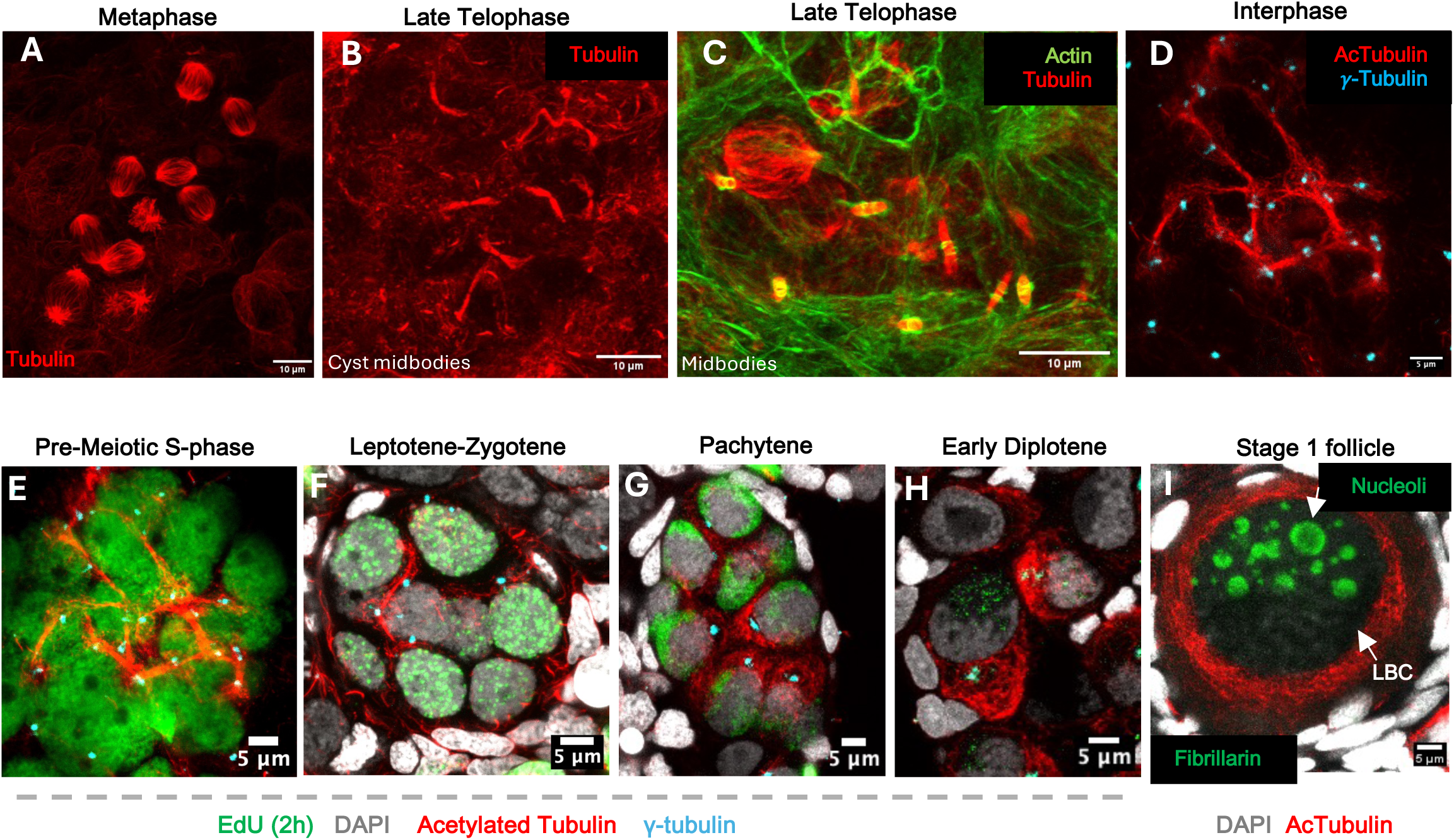
FLS during germline cyst development. **A** – Two germline cysts (an 8-cell cyst and a nearby 2-cell cyst) in metaphase stained with α-tubulin (red). **B** – A 16-cell cyst during late telophase showing typical 8 midbodies stained with α-tubulin (red). **C** – 8-cell cyst with transforming midbodies to more elongated without gaps, spanning intercellular bridges. **D** – During interphase, oogonial cysts form fusome-like structures (FLS) interconnecting sister cells via microtubules. Ovary stained for acetylated microtubules (red) and γ-tubulin (centrioles, cyan). **E–H** – Whole juvenile Xenopus ovaries stained for acetylated microtubules (red), γ-tubulin (cyan), EdU (green), and DAPI (gray). All figures shown as a maximum projection of selected optical slices. **E** – Pre-meiotic S-phase cyst marked by EdU (green), with FLS microtubules (acetylated MT, red) extending from MTOCs (γ-tubulin, cyan) in each cell. **F** – Leptotene–zygotene stage cyst showing reduced FLS structure. **G** – Pachytene-stage cyst with further reduced FLS. **H** – Early diplotene primordial follicle recently separated from the cyst, with microtubules accumulating at the vegetal pole around the forming Balbiani body. **I** – Stage I follicle showing cytoplasmic and nuclear growth and a germinal vesicle (GV) with multiple nucleoli (fibrillarin, green), each associated with extrachromosomal rDNA copies, and lampbrush chromosomes (LBC). Microtubules are enriched around the GV.

Early meiotic prophase occurs within vertebrate and invertebrate germline cysts suggesting that cyst formation evolved as an animal prelude to meiotic entry. In Drosophila, stage-specific changes in the fusome have long been noted during meiosis. Consequently, we next studied the Xenopus FLS during early meiotic prophase (Figure 4E-I, Suppl. Figure S6). Figure 4E-I summarizes changes in the morphology of the FLS during an approximately 30-day developmental period between pre-meiotic S phase and diplotene follicles where the cyst has broken down. As in premeiotic cysts (Figure 4E) the FLS in leptotene-zygotene (Figure 4F) continue to span cyst cells, and associate with centrioles. However, the bundles connecting cells become less compact and spread more widely in the cell during pachytene (Figure 4G). During early diplotene (Figure 4H), cysts split into individual primordial follicles, where MTs have spread through much of the cytoplasm and become enriched around the forming Bb on a vegetal pole. Figure 4I illustrates a growing stage I follicle, where chromosomes transform into lampbrush chromosomes, and nucleoli form around each amplified rDNA copy.

### Transport of vesicles and materials between cyst cells via ring canals

We carried out electron microscopy on Xenopus ovaries and analyzed serial ultrathin sections to identify intercellular transport by visualizing organelles within the lumen of ring canals (Figure 5A, B; Suppl. Figure S7). Electron microscopy revealed that intercellular bridges between cyst cells were invariably filled with cytoplasmic material, including Golgi vesicle-like (Figure 5A) and ER-like structures (Figure 5B). Immunostaining for GM130 (Golgi matrix protein) in whole ovaries detected vesicle-like structures colocalized with microtubules crossing the bridge (Figure 5A’, A”). Similarly, immunostaining for PDI revealed ER structures associated with acetylated microtubules in bridges between oogonia cystocytes (Figure 5B’, B”).

**Figure 5.**
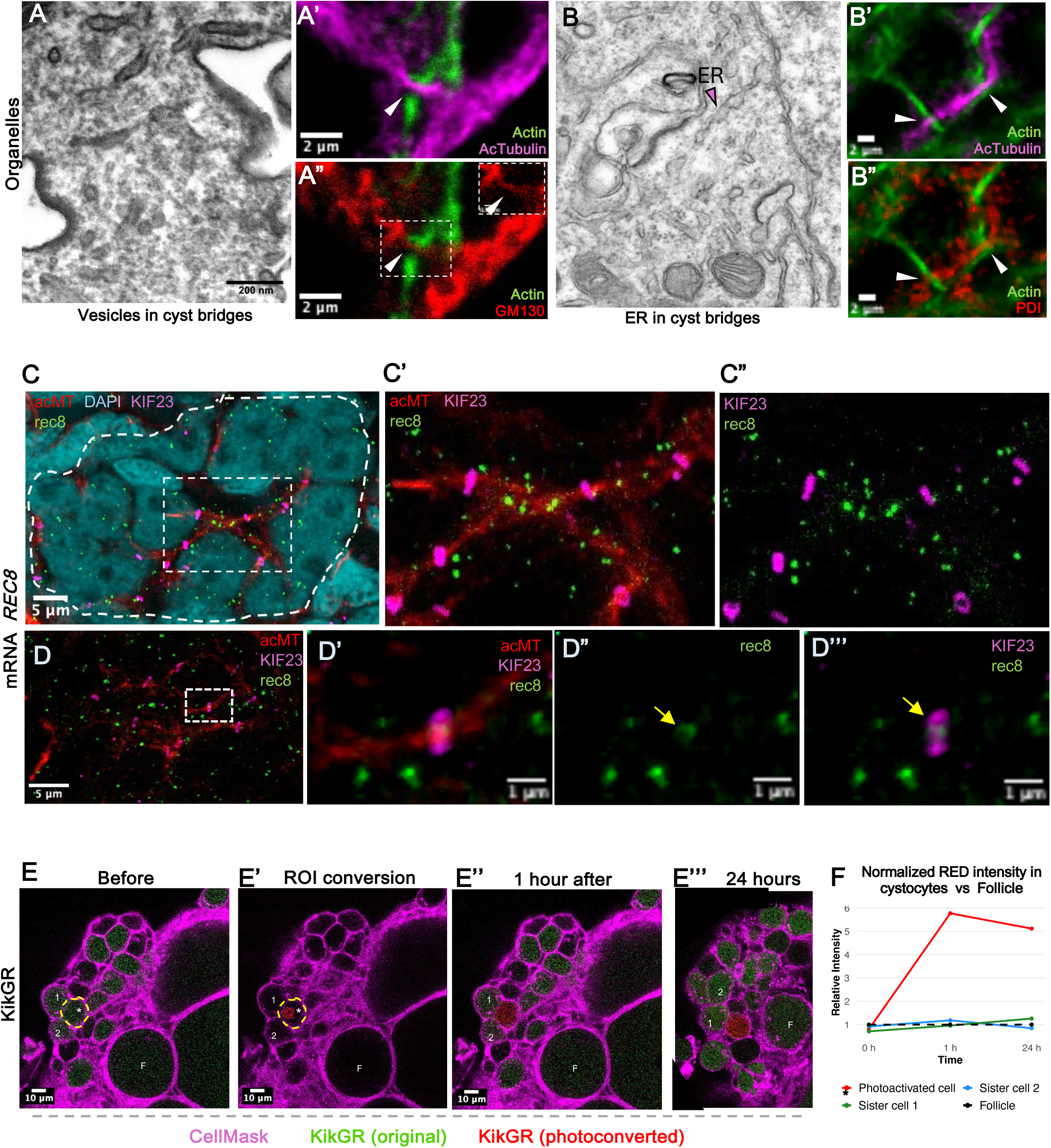
Vesicles, ER, and mRNA – but not photoconvertible KikGR – are shared through ring canals. **A–A’’** – Golgi-derived vesicles are observed within intercellular bridges (IBs). **A** – Electron microscopy of an IB shows multiple vesicles within its lumen. **A’–A’’** – Immunofluorescence (IF) of whole-mount ovary stained for acetylated tubulin (magenta), phalloidin (green), and GM130 (red) reveals Golgi-derived vesicles localized along microtubules (arrowheads). **B–B’’** – ER-like structures are present within intercellular bridges. **B’–B’’** – IF of whole-mount ovary stained for acetylated tubulin (magenta), phalloidin (green), and PDI (red) shows colocalization of ER with microtubules crossing intercellular bridges (arrowheads). **C–C’’** – HCR RNA-FISH with a *rec8* probe (green), combined with IF for acetylated tubulin (red), ring canals (Kif23, magenta), and DAPI (cyan). **C’–C’’** – Zoom-in from the boxed region in C showing *rec8* mRNA enriched on microtubules in a cystocyte with four ring canals. **D–D’’’** – HCR RNA-FISH with *rec8* probe (green), combined with IF for acetylated tubulin (red), ring canals (Kif23, magenta), and DAPI (cyan). **D’–D’’’** – Zoom-in from the boxed region in D showing individual *rec8* RNA molecule located within the lumen of ring canal. **E–E’’’** – Photoconversion of a single cyst cell in a dissected ovary from KikGR green to red. Cell membranes are labeled with CellMask (magenta). The converted cell is outlined in yellow and marked with an asterisk. **E** – Before conversion. **E’** – ROI showing conversion of a single cell. **E’’** – 1 hour post-conversion. **E’’’** – 24 hours post-conversion. **F** – Quantification of red signal intensity in neighboring cystocytes versus an independent follicle not connected to the cyst. Y-axis: relative red intensity; X-axis: time. Sister cells 1 and 2 correspond to those indicated in E–E’’’; follicle (F) used for normalization.

We also examined how various organelles change in a cyst during meiotic stages (Suppl. Figure S8). During oogonial stages ER can be found co-locolized with microtubule bundles. ER appears enriched in a subregion of cyst cells that contains the centrosome and Golgi (lower part of each panel). In the early diplotene stage shown, the cyst has broken down entirely and has formed a new primordial follicle with ER enrichment around the forming Bb. Golgi and γ-tubulin staining shows that centrosomes and Golgi are also located asymmetrically in the cell (Suppl. Figure S8B). In Drosophila, centrioles migrate through intercellular bridges along the fusome toward the pro-oocyte in late pachytene. In mouse nurse cell centrosomes, mitochondria, Golgi and cytoplasm pass through membrane gaps and into the oocyte (Lei and Spradling, 2016). To investigate whether directional organelle transfer occurs in Xenopus, we stained centrioles using γ-tubulin and centrin-2 (Suppl. Figure S8C-D). γ-Tubulin foci expanded at the pachytene stage and declined at the stage 1 follicle, but it was unclear whether these changes were associated only with centrioles. In contrast, centrin-2 consistently marked two distinct dots per cystocyte, indicating that each cystocyte retains its own centrioles. No evidence of centriole transfer was observed based on constancy in the number of centrin-2 foci within individual cells. Both electron microscopy and immunofluorescence failed to detect mitochondrial passage through Xenopus intercellular bridges (data not shown). Ring canal diameters at different cyst stages remained relatively constant throughout all stages (Suppl. Figure S9).

To test whether RNA passes through ring canals, we performed hybridization chain reaction FISH (HCR-FISH) for *rec8*, a marker of premeiotic S phase, combined with immunostaining for Kif23 and acetylated tubulin. *Rec8* mRNA localized along microtubule bundles, particularly in cystocytes with multiple ring canals (Figure 5C-C’’), and was occasionally detected within the center of ring canal lumens (Figure 5D-D’’’), suggesting mRNA transport between cyst cells. Whether this transport is directional or selective for specific transcripts, such as oskar mRNA in Drosophila, remains an open question. To test whether cystocytes exchange cytoplasmic proteins, we performed live imaging of ovaries from transgenic frogs (*Xla.Tg(CAG:KikGR*). The KikGR (green) protein undergoes conversion to red fluorescent protein following photoconversion using UV light (Tandon et al., 2013). Isolated ovaries were stained with CellMask™ plasma membrane dye, and KikGR protein was photoconverted from green to red in a single cystocyte using UV light. Imaging before and after photoconversion (1 hr and 24 hr post conversion) showed no detectable transfer of labeled proteins between cystocytes (Figure 5E-F). Neither green KikGR protein entered the photoconverted cell, nor did red KikGR protein diffuse to neighboring cystocytes. These findings suggest that intercellular bridges in *Xenopus* do not facilitate significant KikGR protein sharing over a 24-hr period.

### Microtubule connections between cystocytes can reform following disruption

To assess the role of microtubule connections in cyst development and intercellular exchange, we examined their recovery following complete depolymerization. While spindle remnants likely initiate intercellular microtubule formation, we tested whether microtubules could reform independently of midbody-derived structures. To test this, we treated ovaries with Nocodazole, which completely depolymerized acetylated microtubules in germline cysts within 15– 20 hours (Figure 6A, B). Notably, in the whole ovary only cells with features of GSCs retained acetylated microtubules even after 20 hours of nocodazole treatment. Importantly, ring canals remained unaffected (Figure 6B), and the vast majority of cysts appeared healthy, although some showed signs of degradation. Following drug washout, cytoplasmic microtubules fully recovered within 10–24 hr. Notably, we also observed microtubule recovery within intercellular bridges at all germline cyst stages (Figure 6C), suggesting that microtubules can regrow within bridges independently of spindle remnants. To avoid the confounding effect of new spindle formation in dividing oogonial cysts during recovery, we focused our analysis on meiotic-stage cysts, which progress more slowly and are unlikely to divide within the recovery time frame. Quantification revealed that prior to treatment, approximately 80–95% of analyzed intercellular bridges in meiotic cysts at leptotene–zygotene and pachytene stages contained microtubules. Following nocodazole washout, a similar percentage of bridges showed microtubule recovery and reconnection (Figure 6D). After microtubule recovery, particularly in pachytene-stage cysts, we frequently observed long MT bundles—up to 20–30 μm in length—spanning between sister cells that had moved apart but remained connected by intercellular bridge (Suppl. Figure S10). Such configuration was never seen in untreated controls, suggesting that in the absence of MTs, cyst cells may drift apart without fully disconnecting.

**Figure 6.**
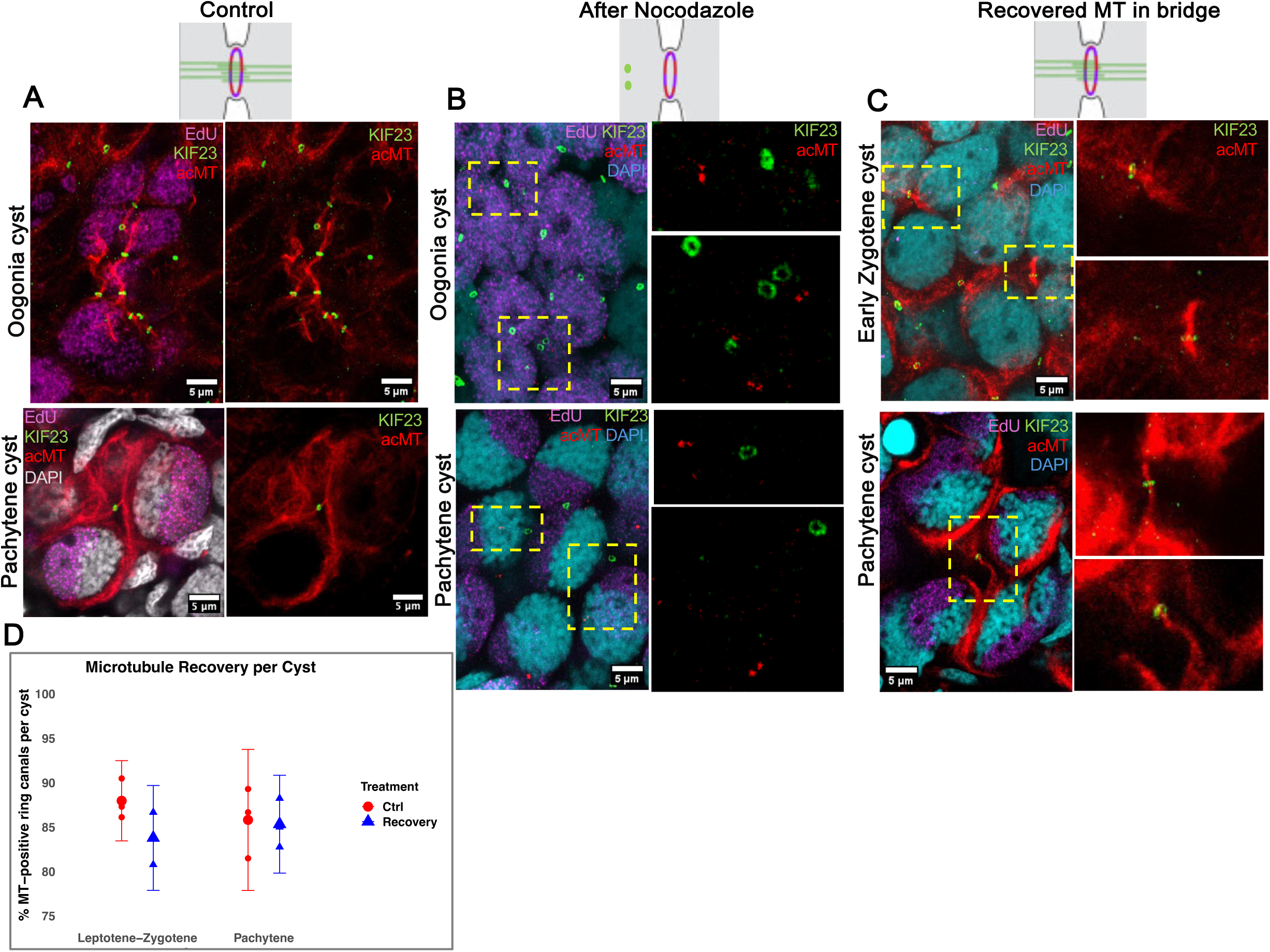
Microtubule interconnections between cystocytes are actively maintained. All samples are stained with acetylated tubulin (red), Kif23 (green), EdU (incubated for 1h before recovery, magenta), and DAPI (gray or cyan). **A** – Control whole juvenile ovaries treated with DMSO for the same duration, showing microtubule networks in oogonia and pachytene cysts. **B** – Ovaries immediately after nocodazole treatment (0 hr), showing depolymerization of microtubules in oogonia and pachytene cysts. Ring canals remain intact, but microtubules are reduced to centrioles. **C** – Ovaries after nocodazole washout and recovery, showing regrowth of microtubules within intercellular bridges in early zygotene and pachytene cysts. **D** – Quantification of microtubule recovery in cysts. The percentage of bridges with visible microtubules was measured for leptotene–zygotene and pachytene cysts in control and recovery conditions. No microtubules were detected in bridges at 0 hr after nocodazole treatment.

### Xenopus cysts contain nurse cells that turn over and do not form oocytes

Previously, all individual cystocytes were proposed to survive and differentiate into oocytes, based on a dearth of TUNEL-positive nuclei observed in the post-metamorphic ovary (Kloc et al., 2004a). Subsequently, studies in Drosophila (review: Lebo and McCall, 2021), and later in mouse (Niu and Spradling, 2022) showed that apoptosis is not used routinely for nurse cell turnover by these species. Instead, a process of programmed cell death involving acidification by somatic cells and degradation near other cyst cells takes place (Mondragon et al., 2019; Lebo and McCall, 2021). We developed a lineage-tracing method using EdU pulse-chase labeling, since a Cre-loxP genetic system is not available for *Xenopus* to revisit this issue.

We injected EdU into 125 metamorphic Xenopus tadpoles and dissected their gonads every two days to track cystocyte fate. This approach enabled us to establish the timeline of cyst development, tracing the progression from oogonial stages through early meiotic prophase (leptotene, zygotene, and pachytene) to the formation of individual follicles. Oogonial cysts appeared 0–4 days after the EdU pulse, leptotene–zygotene cysts at 2–9 days, late zygotene-early pachytene cysts at 9–15 days, late pachytene cysts at 11–18 days, early diplotene cysts at 15–27 days, and stage I follicles after 27 days (Suppl. Figure S11). These findings suggest that, on average, cystocytes require approximately one month to progress from oogonial divisions and early meiotic prophase to the formation of individualized stage 1 follicles which coincides with a conclusion from Coggins and Gall, 1972.

Because our labeling method only marks cysts undergoing S-phase at the time of EdU exposure, it allows a straightforward situation where cell turnover can be followed over subsequent days with comparative quantification of developmental stages (Figure 7A, B, D). It is clear that over this 30 day period, the great majority of cell labeling is lost. We estimated from sampling that there were at least 1,296 germ cells labeled 4d after the EdU pulse. At 33 days, the remaining signal originates from compact somatic nuclei dispersed throughout the ovary, whereas germ cells at this stage are organized into follicles with developing lampbrush chromosomes, which are only visible upon zooming in (Figure 7B, rectangle box). Moreover, it was easy to count the 217 EdU-labeled cells remaining at 33 days, which microscopic examination verified were all no longer cysts, but young ovarian follicles (Figure 7B, rectangle box; 7D). Thus, it is clear that the great majority of cyst cells (∼83%) do not survive to become oocytes (Figure 7D).

**Figure 7.**
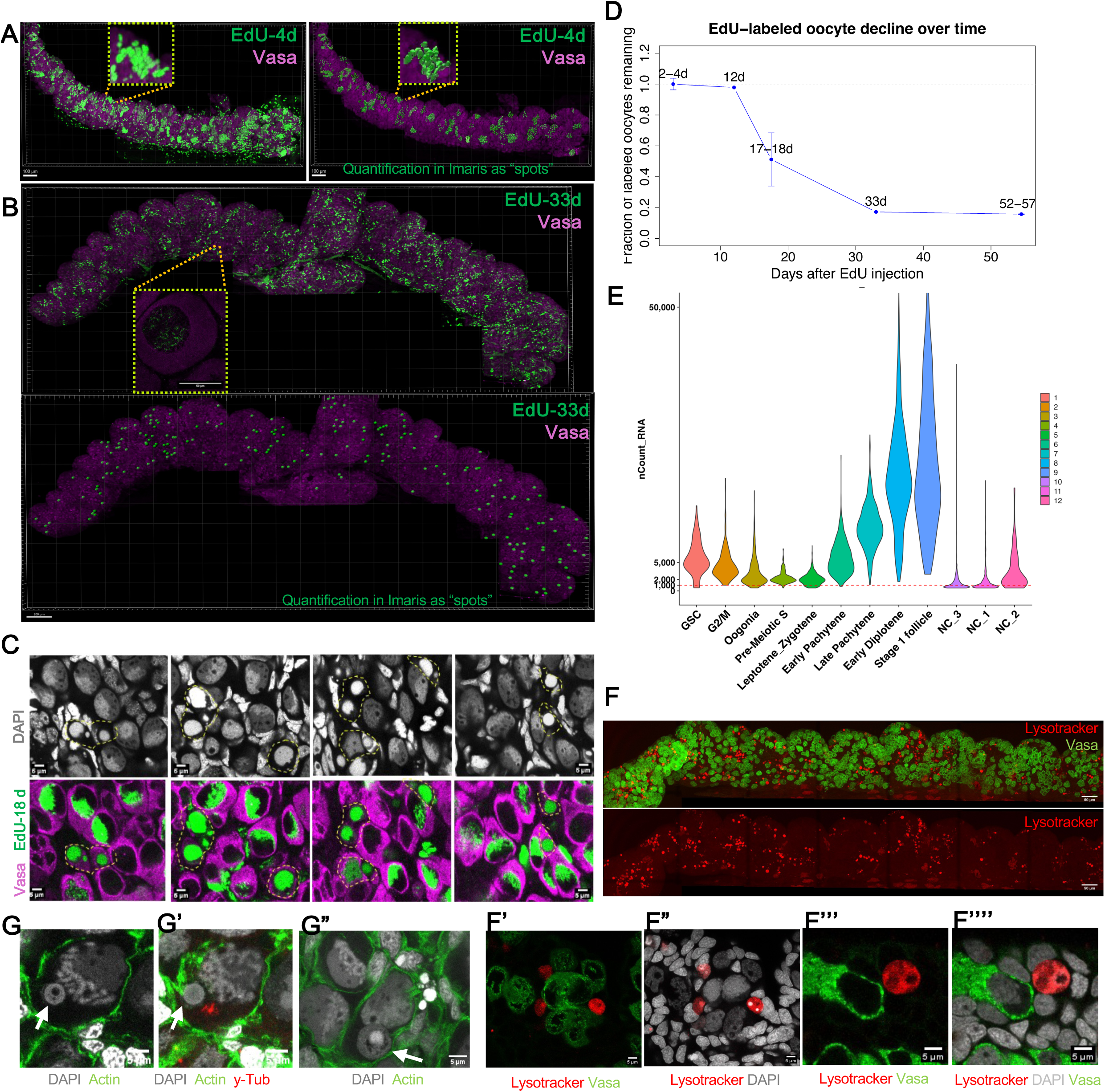
Most cyst cells are putative nurse cells that undergo turnover. **A–A’** – Whole juvenile ovary 4 days after EdU injection. Germline cells (Vasa, magenta); EdU (green). **A’** – Quantification performed manually in Imaris using the ‘Spot’ and ‘Cell’ functions. Dashed boxes indicate germline cysts shown in original (A) and reconstructed views (A’). **B–B’** – Same staining and analysis as in A, shown for an ovary 33 days after EdU injection. **C** – Dying EdU-positive germline cells observed during late pachytene/early diplotene stages as cysts fragment (EdU, magenta; Vasa, green; DAPI, gray); whole ovary at 18 days post-EdU injection. **D** – Quantification of the fraction of EdU-labeled oocytes remaining in the ovary from 2 to 57 days after EdU pulse. **E** – Distribution of nCount_RNA (UMI counts) per cell across indicated germline clusters. **F** – Extensive nurse cell turnover by programmed cell death visualized in juvenile ovaries. Acidified nuclear remnants (Lysotracker, red); germline cells (Vasa, green). **F’–F’’’’** – Zoomed regions from F showing germline cells adjacent to acidified nuclear remnants. **G** – Nuclear remnants from dying nurse-like cells are sometimes seen within early pachytene oocytes. Staining: actin (green), DAPI (gray), γ-tubulin (red).

Since we examined samples at several intermediate time points, we observed that relatively little turnover took place early, but significant amounts of EdU-positive germ cell death began around 18 days. At this time, most EdU-labeled cystocytes were in late pachytene-early diplotene stages, and many EdU-positive germ cells nearby exhibited morphological features consistent with cell death (Figure 7C). These included a smooth nucleus lacking the characteristic chromosomal bouquet and rDNA cap, as well as a shrunken, Vasa-positive cytoplasm. Such a phenotype of germ cell death was also observed in ovaries without EdU pulse-chase which proves that it’s not an artefact of cell death due to EdU incorporation.

Moreover, as was mentioned earlier, scRNAseq data revealed clusters in the inner ring with relatively low RNA counts per cell (UMI) (Figure 1A, 7E). One cluster (#11 or NC1) is positioned near leptotene/zygotene cells near the start of meiosis and near early diplotene cells when cyst tends to break down, while another cluster (#10 or NC3) is close to the stage 1 follicle which could represent rather wave 1 follicle death (Niu, Spradling 2020; Yin, Spradling 2025). Together, these clusters (#11 or NC1, #10 or NC3, #12 or NC2) account for about 20% of the germline cells (or 10% of NC1 from total germline cells), which represents a substantial fraction of cells, given the short lifetime of nurse cells compared to oocyte precursors.

To determine the cell death pathway—whether it occurs through programmed cell death via acidification, as shown in flies (Mondragon et al., 2019; reviewed in Lebo and McCall, 2021) and mice (Niu and Spradling, 2022)—we performed Lysotracker staining on *Xenopus* ovaries. Our results showed that the majority of nuclear remnants were Lysotracker positive (Figure 7F-F’’’’), consistent with previous observations on low apoptotic cell death in the Xenopus ovary (Kloc et al., 2004). Occasionally, similar nuclear remnants were observed within or nearby early diplotene oocytes (Figure 7G–G’’), possibly representing the remnants of degrading nurse cells.

## DISCUSSION

### Transcriptional identity of germline stem cells in Xenopus

Our single-cell RNA sequencing atlas of Xenopus early germline development provides the first comprehensive transcriptional roadmap spanning quiescent germline stem cells (GSCs), mitotically dividing cysts, early meiotic cells, and primordial follicles. For the first time, we were able to capture transcriptomes of rare GSCs and validate their identity by *piwil4* expression using RNA FISH. These GSCs exhibit a distinct transcriptional profile characterized by hallmark features of vertebrate germline stemness: globally low translation, activation of JAK1/STAT1 signaling, and expression of transposon-silencing effectors (*piwil4, spocd1, morc1, dnmt3a*).

The molecular identity of *Xenopus* GSCs reveals additional evolutionary parallel with germline stem cells in other vertebrates. For instance, they share a neuronal-like gene signature— including *chgb*, *sncb* (synuclein beta, typically found in presynaptic terminals), and the neuropeptide *npy*—with zebrafish GSCs (Liu et al., 2022). They also express *piwil4* (the ortholog of mouse *Miwi2*), which is a proposed marker of early spermatogonia in the human testis (Kwaspen et al., 2024), as well as core components of the piRNA pathway involved in transposon repression during fetal male germline development in mice (Uneme et al., 2024).

Notably, *Xenopus* GSCs express *tubb3* (class III β-tubulin), an isoform previously reported only in terminally differentiated neurons (Liu et al., 2007) and Sertoli cells (Gendt et al., 2011). In those contexts, *tubb3* is regulated by androgens, which influence cytoskeletal gene expression and promote resistance to cellular stress (Butler et al., 2001; Gendt et al., 2011). Strikingly, *Xenopus* GSCs were the only ovarian cell type that maintained highly stable, acetylated microtubules under nocodazole treatment, suggesting enhanced microtubule resilience. These findings raise the possibility that hormonal signals—perhaps androgen-like—may contribute to cytoskeletal stability and GSC maintenance in amphibians.

The fact that adult Xenopus females can be induced to ovulate multiple times per year (Moss et al., 2024) and are capable of regenerating new ovarian lobes after partial ovariectomy, supports the conclusion that Xenopus GSCs are not merely rudimentary remnants but are functionally competent stem cells with the capacity to regenerate new cysts and follicles throughout adult life.

### Xenopus oocytes arise in germline cysts with a prominent fusome-like structure

Xenopus ovarian cysts arise with many of the features that have been described previously in male gametogenesis and most of the characteristics of female premeiotic cells from a wide diversity of animal groups. These include mitotic synchrony, incomplete cytokinesis, formation of a stabilized intercellular bridge, assembly of a microtubule-rich fusome from spindle remnants, readjustment of bridges toward each other in a rosette pattern, and asymmetric accumulation of fusome microtubules and associated materials preferentially in cyst cells containing multiple branches. Two kinds of cell derive from Xenopus cysts, a relatively small number of oocytes, and a large number, 80% of more, of nurse cells defined as cyst cells that are programmed to serve functions other than that of oocytes. No longer can lower vertebrates be considered an exception to the evolutionary conservation represented by the cyst program.

A key feature of the Xenopus FLS is the exceptional strength and stability of its microtubule-rich connections. These MTs span cystocytes throughout interphase, peak during premeiotic S-phase, and gradually disperse as meiosis progresses—mirroring the developmental pattern of Drosophila fusomes (Grieder et al., 2000). Our observations suggest that this structural stability may be essential for maintaining cyst architecture and coordinated development.

Importantly, the FLS demonstrated remarkable plasticity: when microtubules were experimentally depolymerized, cystocytes could drift apart from each other yet remained tethered by ring canals. Upon recovery, long microtubule bundles regrew, spanning even distant cells. Our experiments support the idea that microtubule connections within a cyst are essential not only for maintaining mechanical cohesion and cyst conformation, but also for enabling molecular sharing among cystocytes.

Although rare, we observed cyst fragmentation during oogonial stages. Live imaging and fixed material revealed that oogonia exhibit increased motility compared to meiotic cysts, as also recently demonstrated in mouse (Levy et al., 2024). This motility likely imposes mechanical stress on intercellular bridges. As far as we can tell, fragmentation tended to occur at the cyst periphery, where fusome microtubules are weaker due to their asymmetric distribution. This suggests that the FLS may play a role in mechanically stabilizing cysts and protecting intercellular bridges from rupture during periods of increased cell motility.

### Future nurse cells downregulate key meiosis genes early

Nurse cells have very different tasks than oocytes, and it has long been known in Drosophila that only the future oocyte and one nurse cell form a synaptonemal complex and prepare for recombination (Carpenter, 1975). Our results suggest analyzing gene changes in NC1 nurse cells show that a similar process takes place in Xenopus cysts. Once specified, nurse cells have nothing to gain by continuing with synapsis and recombination, since their genome will never be used in a gamete. Instead, the true oocyte stands to benefit by ensuring that supportive nurse cell transcriptional will not be adversely affected by ongoing recombination, replication, DNA repair and meiotic checkpoint activation that could reduce transcriptional output and inactivate chromatin in some chromosome regions.

We examined the earliest Xenopus nurse cells, NC1, which appear in our scRNAseq profile at early leptotene. NC1 nurse cells express a suite of genes that can be understand as a program for nurse cell development from an uncommitted cyst cell that retains oocyte potential. They downregulated genes controlling the chromosomal aspects of meiosis, including Sycp1, hormad1, and Dmc1 all of which are required for homologous chromosome synapsis in oocytes and mutate to sterility. Synapsis in turn is essential for recombination and chiasma formation. Hormad1 promotes synapsis by binding to unpaired sites, and initiates silencing of genes in unpaired zones. Downregulating these genes will result in what has been observed in Drosophila nurse cells, little or now initial synaptonemal complex formation and an apparent withdrawal from subsequent aspects of the meiotic cycle, while maintaining high levels of gene expression. Our observation that largely the same genes are downregulated at the onset of Xenopus and mouse nurse cell formation marks this as a nurse cell developmental program that may be ancient and conserved in many animal groups. A list of the genes down regulated more than 3-fold in cluster NC1 is given in Table S3 and also includes genes that modulate the the cytoskeleton, translation, chromatin and endocytic trafficking.

The nature and value of such a program can explain a longstanding observations in the study of cysts that has not been well understood. It has been widely observed for >100 years cysts formed by synchronous divisions produce only one oocyte and 2n-1 nurse cells, a fact known as the “2n rule” (Giardina, 1901; Buning, 1994). The realization that nurse cells abort meiotic chromosome behavior for likely benefit can explain the 2^n^ rule, and explains why early cyst breakage of a polarized structure can to smaller cysts that each generate one oocyte, as seen in mouse oogenesis (Lei and Spradling, 2016).

### Xenopus cysts produce oocytes and cells with some but not all nurse cell features

The vast majority of insect species, like Drosophila, produce cysts with a majority of nurse cells that transfer materials to the oocyte before turning over. In mammals such as mouse, about 80% of cyst cells turn over within cysts during pre-follicular oogenesis after transferring organelles and cytoplasm to oocytes (Lei and Spradling, 2016; Niu and Spradling, 2022). Here we succeeded in showing by EdU-marking that ∼80% of Xenopus cyst cells also turn over prior to follicle formation. This dichotomy of cell fates was also clearly revealed by scRNA sequencing. The Xenopus profile showed a similar picture as in mouse, in which about 10% of low UMI nurse-like cells from cluster NC1 were found on the UMAP plot inside of the main sequence meiotic cells. These cells are probably transferring some materials through FLS since the growth of UMI in Xenopus meiotic cells with stage was very similar to mouse, and shrunken acidified germ cells typical of dying nurse cells in Drosophila and mouse were observed. Clearly, lower vertebrates are not a general exception to the germline cyst paradigm by producing only oocytes (Kloc et al., 2004a; Brubacher, 2024).

Xenopus nurse-like cells almost certainly carry out valuable functions prior to their turnover. Our findings suggest that nurse-like cells may participate in the selective transfer of cytoplasmic materials, including Golgi-derived vesicles, ER, and mRNA, rather than bulk cytoplasm including organelles, possibly via fusome-associated transport mechanisms. They may transfer key protective molecules such as small RNAs and lncRNAs, as suggested by the experiments looking at transfer of *rec8* mRNA. Additionally, our experiments with the photoconvertible protein KikGR, which lacks cellular function, suggest that non-essential proteins are not shared through intercellular bridges. Furthermore, we did not detect transfer of organelles in bulk like in mouse or Drosophila cysts: ring canal diameters did not increase with development, organelles did not accumulate disproportionately in any one cystocyte, and centrioles (marked by centrin-2), remained equally distributed. These results suggest that the oocyte in Xenopus may not rely on early bulk organelle transfer within the cyst but instead amplifies organelles during the later lampbrush chromosome stage, when transcription is extremely active (Gall, 2012).

This strategy contrasts with models from Drosophila and mouse, where nurse cells dump large quantities of rejuvenated cytoplasm and organelles to the oocyte during meiotic prophase in cyst. During meiosis in yeast and Drosophila proteostasis enhancement and organelle competitive selection and repair are thought to help generate and identify highly functional organelles (Cox and Spradling, 2003; Unal et al. 2010; Goodman et al. 2020; Palozzi et al. 2022; Spradling et al. 2022; Monteiro et al. 2023). The rejuvenated organelles are then deposited into the oocyte, where they support early embryogenesis. Accordingly, Balbiani bodies in mouse and Drosophila are transient, dispersing rapidly after organelle transfer. In contrast, Xenopus and other lower vertebrates may take a delayed approach: organelles are not rejuvenated during the cyst stage, and bulk transfer is absent. Instead, the oocyte may gradually improve organelle quality during its long diplotene stage, during which the Balbiani body persists (Tworzydlo et al. 2016). This may explain why the Balbiani body is longer-lasting in Xenopus, possibly allowing time for internal organelle selection and maturation.

A related but distinct strategy is observed in some marsupial frogs, where hundreds to thousands of transcriptionally active nuclei co-exist within a single oocyte (del Pino, 2018, 2020). These nuclei contribute biosynthetically—producing RNAs and proteins to support a shared cytoplasm. Eventually, only one nucleus is retained for meiosis, and the others degrade. Although structurally different, this mirrors the Xenopus system in which ∼80% of cystocytes are eliminated, but may transiently serve a nurse-like role. Together, these examples suggest that the key evolutionary function of nurse cells may not always be bulk organelle transfer. Rather, nurse cells—or nurse-like cells—may broadly serve to biosynthetically support the oocyte, either by contributing organelles, transcripts, or cytoplasmic components, depending on the species. The balance between these strategies likely reflects developmental constraints and evolutionary adaptations specific to each lineage.

## METHODS

### Experimental model and ethical approval

The experimental model in this study is *Xenopus laevis*. Xenopus tadpoles with 4 limbs (NF59-62), late metamorphic froglets (NF63-65) or sexually mature females were purchased from Xenopus1 company (https://xenopus1.com/). All experiments in this study were performed in accordance with protocols approved by the Institutional Animal Care and Use Committee (IACUC) of the Carnegie Institution of Washington.

### Whole-mount staining of Xenopus ovaries

Whole gonads were dissected under a stereomicroscope, briefly washed in L-15 medium, and immediately fixed in freshly prepared 4% paraformaldehyde (PFA) in PBS for 1.5 hr or overnight at room temperature (RT) based on the experimental requirements. Post fixation, tissues were washed in PBS and permeabilized in PBST (PBS with 0.5% Triton X-100 and 0.1% Tween-20) for 1 hr then blocked in 10% BSA in PBST for 1 hr at RT. Tissues were incubated with primary antibodies for 2 days at RT with mixing, followed by thorough washing in PBST for 1.5 hr. Secondary antibodies, DAPI (1 µg/ml), and, if required, Alexa 488-Phalloidin (Thermo Fisher, 1 µg/ml) to visualize cell membranes were applied overnight at RT. After a final 1.5 hr wash in PBST, samples were mounted in Vectashield® vibrance antifade mounting medium and imaged using a Leica DIVE confocal microscope.

### List of primary antibodies

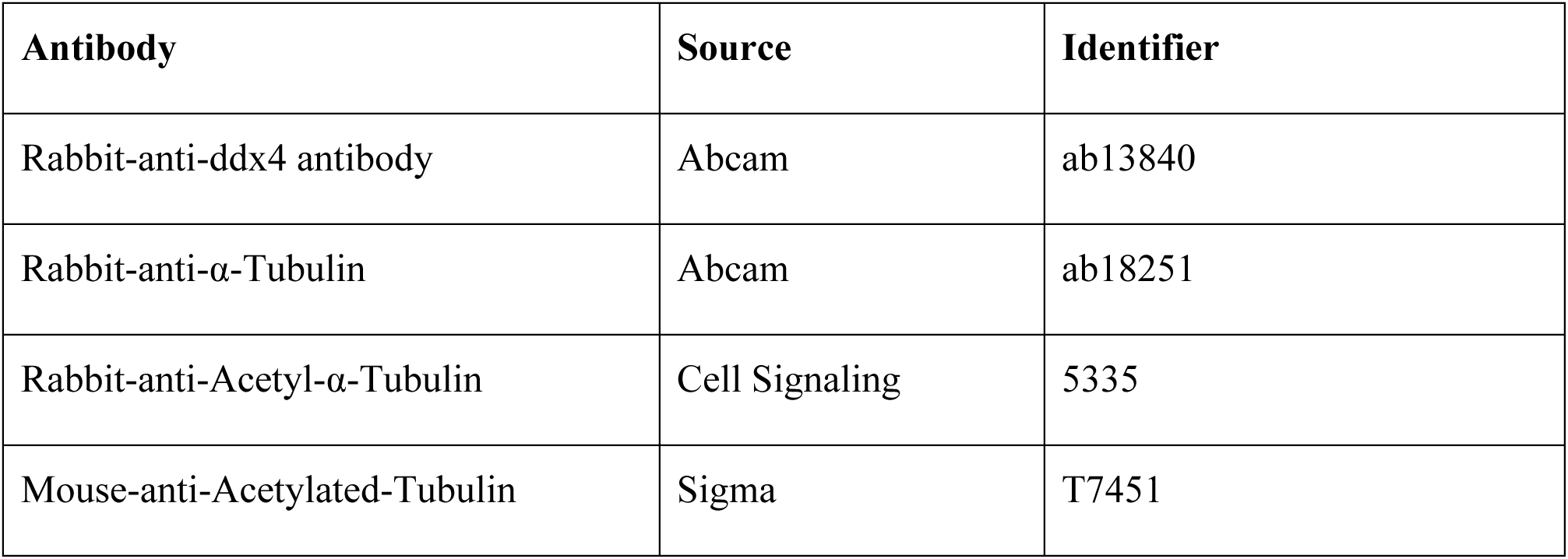
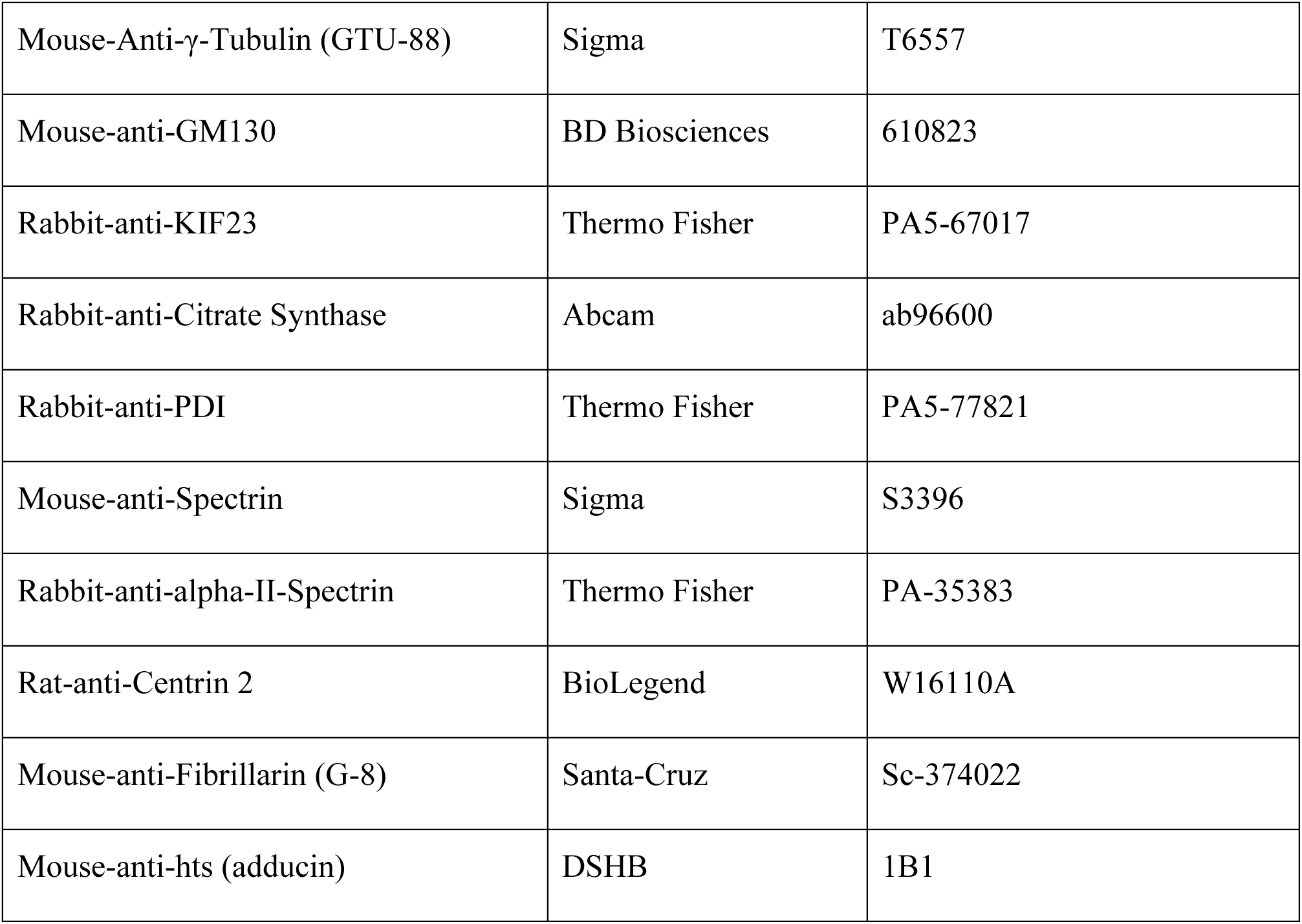

| **Antibody** | **Source** | **Identifier** |
| --- | --- | --- |
| Rabbit-anti-ddx4 antibody | Abcam | ab13840 |
| Rabbit-anti-α-Tubulin | Abcam | ab18251 |
| Rabbit-anti-Acetyl-α-Tubulin | Cell Signaling | 5335 |
| Mouse-anti-Acetylated-Tubulin | Sigma | T7451 |
| Mouse-Anti-γ-Tubulin (GTU-88) | Sigma | T6557 |
| Mouse-anti-GM130 | BD Biosciences | 610823 |
| Rabbit-anti-KIF23 | Thermo Fisher | PA5-67017 |
| Rabbit-anti-Citrate Synthase | Abcam | ab96600 |
| Rabbit-anti-PDI | Thermo Fisher | PA5-77821 |
| Mouse-anti-Spectrin | Sigma | S3396 |
| Rabbit-anti-alpha-II-Spectrin | Thermo Fisher | PA-35383 |
| Rat-anti-Centrin 2 | BioLegend | W16110A |
| Mouse-anti-Fibrillarin (G-8) | Santa-Cruz | Sc-374022 |
| Mouse-anti-hts (adducin) | DSHB | 1B1 |

### LysoTracker staining

Dissected ovaries were cultured in vitro in 1 mL of 50 nM LysoTracker Red (L7528, Thermo Fisher) diluted in L-15 medium supplemented with 5% FBS and 1× Penicillin-Streptomycin, at room temperature for 6 or 15 hr. Ovaries were then fixed in 4% PFA/PBS for further analysis.

### Live Imaging

For live imaging, to stain nuclei, juvenile ovaries were incubated in L-15 medium with Hoechst 33342 (1:200; ThermoFisher) overnight at RT. The following day, ovaries were rinsed in fresh L-15 medium and mounted in 0.5–1% low-melting agarose in a 35 mm glass-bottom dish (Cat#: D35-20-1.5H) for imaging. After solidification of the agarose, L-15 medium supplemented with 5% FBS and 1× Penicillin-Streptomycin was added to maintain tissue viability during imaging. Z-stack images were acquired using a Leica DIVE confocal microscope with 405 nm excitation, typically using a 40× or 63× oil objective. Ovaries were imaged live within 6 hr of staining.

### Photoconversion Assay and Live Imaging

To assess cytoplasmic protein exchange between cystocytes, we used juvenile Xenopus laevis ovaries from transgenic frogs expressing the photoconvertible protein KikGR [*Xla.Tg(CAG:KikGR)*] (Tandon et al., 2013). Transgenic froglets (stages NF55–66) were obtained from the National Xenopus Resource (NXR). KikGR fluoresces green under standard excitation and irreversibly converts to red fluorescence upon exposure to UV light. Freshly dissected ovaries were stained overnight at RT with CellMask™ Deep Red plasma membrane dye (#C10046, Thermo Fisher) at 1× concentration in L-15 medium to visualize cell boundaries. Ovaries were then transferred to a 35 mm glass-bottom dish with a 20 mm microwell and embedded in 0.5–1% low-melting agarose to allow orientation. Once the agarose solidified, L-15 medium supplemented with 5% FBS and 1× Penicillin-Streptomycin was added to cover the tissue. Germline cysts were identified based on CellMask membrane staining. An ROI (region of interest) within a single cystocyte was selected in a cyst and photoconverted using 405 nm UV laser and Z-stack images were acquired for 1 hr and 24 hr post-conversion using 488 nm, 561 nm, and 647 nm lasers to visualize unconverted and converted KikGR fluorescence and cell membranes, respectively. Z-stacks were captured using a 63× oil immersion objective. At least six photoconversion events were analyzed across multiple ovaries, with consistent results.

Using Fiji, selected z-stacks were processed with Average Intensity Projection to generate a 2D image of each cyst. We then measured the Integrated Density (total intensity) within the region of interest (ROI), adjacent connected cystocytes, and nearby non-cyst cells (follicle). The Integrated Densities of the ROI and neighboring cystocytes were normalized to that of the non-cyst cell/follicle (Figure 5E; asterisk marks the ROI, 1–2 indicate neighboring cystocytes, F marks a nearby follicle).

### EdU and OPP labeling of ovaries *in vitro*

EdU (5-ethynyl-2’-deoxyuridine; Thermo Fisher) was added to freshly dissected ovaries at 1:500 dilution from stock into L-15 medium supplemented with 5% FBS and 1× Penicillin-Streptomycin. To detect ongoing protein synthesis, O-propargyl-puromycin (OPP; Vector Laboratories) was added at a final concentration of 20 µM. Ovaries were incubated with EdU for at least 1 hr at RT or overnight when needed, and with OPP for 2 hr at RT. Following incubation, gonads were fixed in 4% PFA in PBS and permeabilized in PBST before proceeding with Click-iT detection as per the manufacturer’s protocol.

### Lineage tracing with EdU labeling

To determine the developmental timeline and fate of germline cysts, we performed an EdU pulse-chase experiment in Xenopus. A total of 125 metamorphic froglets (NF62-65, includes males and females) were injected with EdU (10 μg per froglet) and sacrificed at two-day intervals for gonad dissection till day 57. Gonads were fixed in 4% PFA in PBS overnight, permeabilized, and processed for EdU detection (ThermoFisher, Cat# C10337) and whole-mount immunofluorescence. To quantify the number of cells per ovary using Imaris, ‘Spots’ function was used to manually count EdU-positive germ cell nuclei. Because the number of ovarian lobes varies between samples, cell counts were normalized to the number of lobes in each gonad (Figure 7D).

### Nocodazole treatment

Dissected ovaries together with kidney were incubated individually in 24-well plate with Nocodazole (1 μg/mL, SML1665, Sigma) in L-15 supplemented with 5% FBS and 1x Penicillin-Streptomycin for 20 hr. Control samples were treated with an equivalent volume of DMSO. Half of each ovary was fixed immediately after drug exposure, considered as 0 hr. The remaining halves were washed every 30 minutes for ∼6 hr in L-15 medium (5% FBS, 1× Pen/Strep) to remove residual drug and then incubated overnight for complete microtubule recovery. The following day, all samples were fixed in 4% PFA in PBS for 1.5 hr at RT. To visualize both microtubules (acetylated tubulin antibody) and ring canals (Kif23 antibody), a 30-minute incubation in detergent solution (1% SDS, 0.5% Tween-20, 50 mM Tris pH 7.4, 1 mM EDTA, 150 mM NaCl) was used following fixation. This treatment enhances ring canal detection, which is otherwise challenging due to their sensitivity to fixation and requirement for antigen retrieval.

### Single-molecule whole mount HCR RNA FISH

Customized HCR RNA-FISH probes for *piwil4.S, nefm.S* and *rec8.L* were obtained from Molecular Instruments. Hybridization was performed according to the manufacturer’s protocol, with the following modifications. Ovaries were fixed in 4% PFA in PBS for 1.5 hr at RT, dehydrated through a series of methanol treatment and stored at –20 °C until used. Rehydrated ovaries were permeabilized for 30 minutes in detergent solution (1% SDS, 0.5% Tween-20, 50 mM Tris pH 7.4, 1 mM EDTA, 150 mM NaCl). Tissues were pre-hybridized and then incubated with probes at a final concentration of 10 nM overnight at 37 °C. After washes and the amplification step, samples were washed in 5× SSCT (5× SSC with 0.1% Tween-20) containing DAPI (1 µg/ml), mounted in Vectashield antifade, and imaged.

### Combined HCR RNA FISH and immunofluorescence

To combine HCR RNA-FISH with immunofluorescence, we implemented the following modifications to the standard protocol. Ovaries were fixed in 4% PFA for 1.5 hr at RT, washed thoroughly in PBST, and incubated in detergent solution for 30 minutes to permeabilize the tissue. Immunostaining was performed using primary antibodies against Kif23 and acetylated tubulin. After secondary antibody incubation and washes, the tissue was fixed in 2% PFA in PBS for 10 min, followed by several PBST washes. Gonads were then either dehydrated through a methanol series or directly processed for HCR RNA-FISH, following the protocol described above.

### Whole-mount telomere DNA FISH

A Cy3-labeled telomere-specific DNA probe (/5Cy3/TTAGGG repeated 7×) was used to label telomeres. After 1.5 hr fixation, samples were permeabilized in PBST for 1 hr, followed by proteinase K treatment (10 µg/ml, 10 min). Tissues were washed, fixed in 2% PFA in PBS for 10 min and rinsed in PBST. Pre-hybridization was performed in buffer containing 50% formamide, 5× SSC, 0.1% Tween-20, and 0.5 mg/ml tRNA for 1 hr or overnight. Probes were diluted to 2–5 ng/µl in hybridization buffer and co-denatured with the sample at 75 °C for 5 min, followed by overnight incubation at 37 °C. Post-hybridization washes were performed stepwise using decreasing concentrations of pre-HB in 2× SSC, then 0.2× SSC and 2× SSC (with 0.1% Tween-20). Samples were stained with DAPI, mounted in Vectashield antifade, and imaged under a confocal microscope.

### Single-Cell ovary dissociation and sequencing

Ovaries from juvenile Xenopus were dissected at developmental stages between NF62–66. Dissections were performed in L-15 medium at RT. A fresh collagenase solution was prepared in L-15 containing 3 mg/ml Collagenase I (Gibco 17018029), 2 mg/ml Collagenase IV (Gibco 17104019), 3 mg/ml Collagenase II (Sigma C6885), and 1 mM CaCl₂. Ovaries were chopped into small pieces in collagenase solution and transferred into low-adhesion 1 ml tubes. Tissues were digested for ∼40 min at RT with rotation. Cells were pelleted at 300 × g for 3 min, washed in PBS, and digested with 1 ml TrypLE Express (Gibco) for 10 min at RT. The reaction was quenched by adding 10% FBS. Cells were washed twice with cold PBS containing 1% BSA, resuspended in 1 ml PBS with 1% BSA, and filtered through 70 µm and then 40 µm cell strainers to remove clumps. Cells were pelleted at 400 × g for 5 min, resuspended in ∼50 µl PBS, and viability was assessed using automated cell counter (Countess™ 3 FL, ThermoFisher). Cell viability was >95% with minimal clumping. Approximately 10,000 live cells per sample were loaded onto the 10X Genomics Chromium system following the manufacturer’s protocol. Libraries were sequenced on an Illumina NextSeq 500 and processed using Cell Ranger 7.2.0 with default settings.

### Single-cell RNA-seq analysis

Single-cell RNA-seq data from three independent *Xenopus laevis* juvenile ovaries were processed using Seurat v5.1.0. Each sample was converted into a Seurat object and merged using CCA-based integration. Low-quality cells were excluded based on gene count (<100 or >8000 genes) and mitochondrial gene content (>1.5%). Gene expression was normalized (LogNormalize method), and variable features were identified (2000 genes, VST method). Data were scaled, principal component analysis (PCA) was performed, and the top 20 PCs were used for clustering (resolution = 2) and UMAP visualization. After integration using IntegrateLayers() with CCA, the dataset was reclustered using the first 20 dimensions of the integrated reduction (resolution = 1) and visualized with UMAP. Germline clusters were selected based on expression of germ cell markers and UMAP position, then subsetted for reanalysis. The germline subset was normalized, scaled, and reanalyzed using PCA, clustering (resolution = 0.5), and UMAP using 30 dimensions of the integrated reduction.

### Electron microscopy

Ovarian tissue was fixed in 2% PFA/2.5% glutaraldehyde and in 0.1 M cacodylate buffer (pH 7.4) at RT for 1–2 hr, then stored at 4°C. Samples were post-fixed in 1% osmium tetroxide, dehydrated through a graded ethanol series, and embedded in epoxy resin. Ultrathin sections (70–90 nm) were cut using an ultramicrotome, collected on copper grids, stained with uranyl acetate and lead citrate, and imaged with a transmission electron microscope.

### Image analysis and 3D reconstruction using Imaris

Imaris software (Bitplane) was used for quantitative and three-dimensional analyses of confocal image stacks. To determine the exact number of cells within germline cysts, the “Cell” module was used to segment nuclei and measure EdU-labeled cells in 3D. The “Spots” function was applied to count EdU-positive nuclei across different time points following EdU injection. For structural visualization of the fusome, the “Surface” module was used to reconstruct acetylated tubulin-labeled fusomes in three dimensions, allowing assessment of their architecture and connectivity within germline cysts.

## ACKNOWLEDGEMENTS

We thank Mahmud Siddiqi for his assistance with optical microscopy and image analysis. We are grateful to Allison Pinder, Joseph Tran, Fredrick Tan, Xiaobin Zheng, Javier Carpinteyro Ponce, and other members of the Carnegie IT staff for their support with scRNA-seq and bioinformatic analysis. We also thank Dr. Ru-Ching Hsia for her help with electron microscopy experiments. We are especially grateful to members of the Spradling laboratory for their valuable feedback on this project and manuscript, and in particular to Ashish Kumar Tiwary for his helpful suggestions.

## DATA AVAILABILITY

The scRNAseq data are in the process of deposition at the NIH GEO database.

