## SupplementalFigures for "Early female germline development in *Xenopus laevis*: stem cells, nurse cells and germline cysts"

### Supplemental Figures

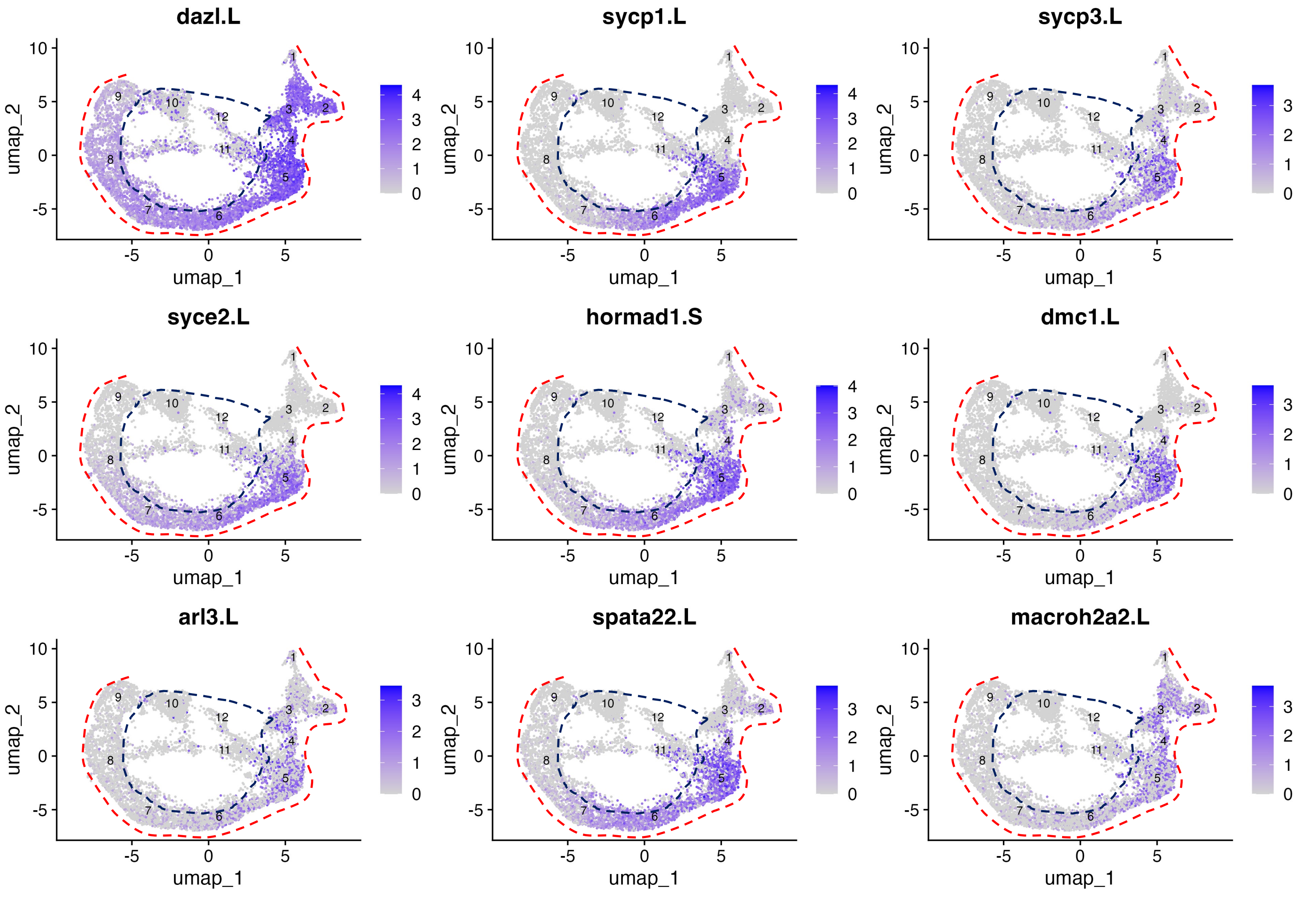

**Figure S1. Genes  downregulated in nurse-like cells compare to meiotic cells in outer ring.**

| Table S3 | Downregulated NC1 nurse cell genes | | | | | | | | |
|---|---|---|---|---|---|---|---|---|---|
| Xenopus | Mouse | LZ exp | NC1 exp | NC1 / LZ | Xenopus | Mouse | LZ exp | NC1 exp | NC1 / LZ |
| faim2 | Faim2 | 7.02 | 0.78 | 0.11 | ano1 | Ano1 | 2.62 | 0.61 | 0.23 |
| sycp3 | Sycp3 | 4.65 | 0.57 | 0.12 | fkbp10 | Fkbp10 | 3.73 | 0.88 | 0.24 |
| dmc1 | Dmc1 | 5.18 | 0.66 | 0.13 | agtrap | Agtrap | 5.75 | 1.44 | 0.25 |
| hormad1 | Hormad1 | 10.11 | 1.50 | 0.15 | dazl | Dazl | 19.69 | 5.01 | 0.25 |
| sycp1 | Sycp1 | 9.68 | 1.61 | 0.17 | c6h7orf57 | | 1.86 | 0.51 | 0.27 |
| macroh2a2 | Macroh2a2 | 3.05 | 0.65 | 0.21 | syce3 | Syce3 | 6.10 | 1.71 | 0.28 |
| syce2 | Syce2 | 6.71 | 1.47 | 0.22 | ccdc63 | Ccdc63 | 1.88 | 0.54 | 0.29 |
| arl3 | Arl3 | 2.51 | 0.55 | 0.22 | hmces | Hmces | 5.32 | 1.50 | 0.29 |
| spata22 | Spata22 | 8.50 | 1.93 | 0.23 | | | | | |
| Downregulated Xenopus genes in NC1 nurse cells compared to leptotene/zygotene (LZ) cells (all genes of 12,154 with NC1/LZ <0.3) | | | | | | | | | |

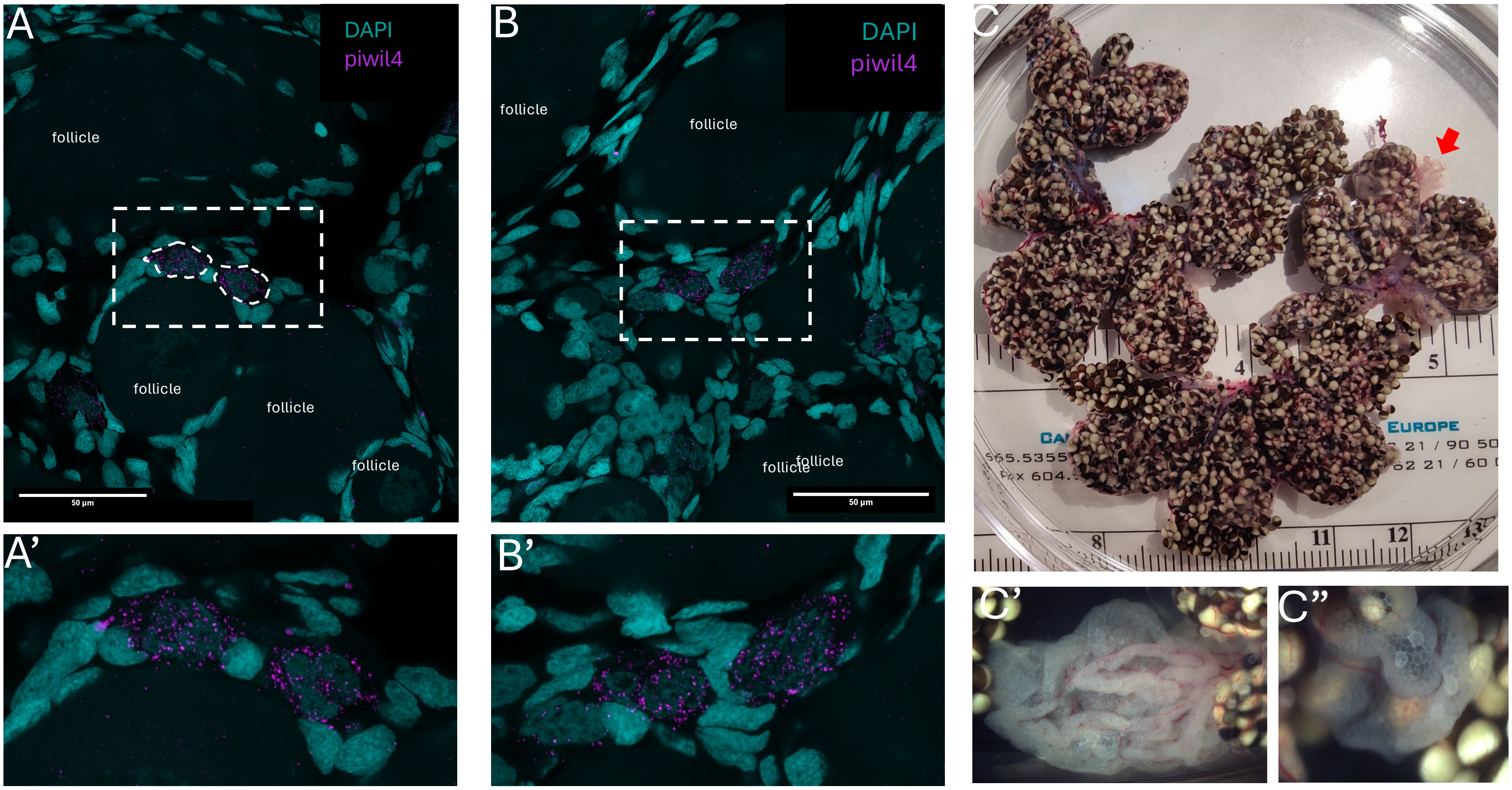

Figure S2. Germline stem cells in the adult  Xenopus ovary.

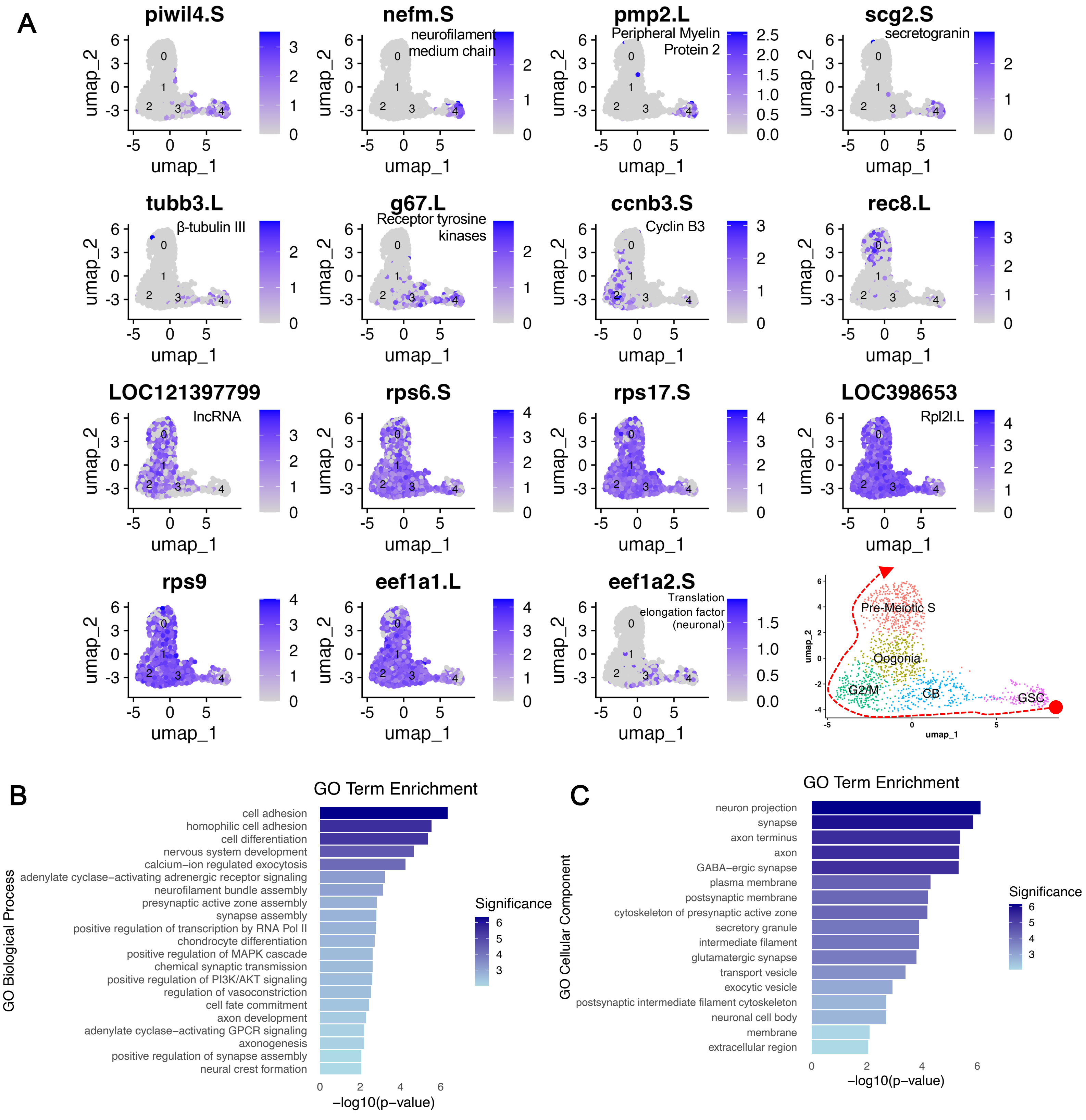

**Figure S3. Expression features of early germline clusters from GSCs to pre-meiotic S-phase.**

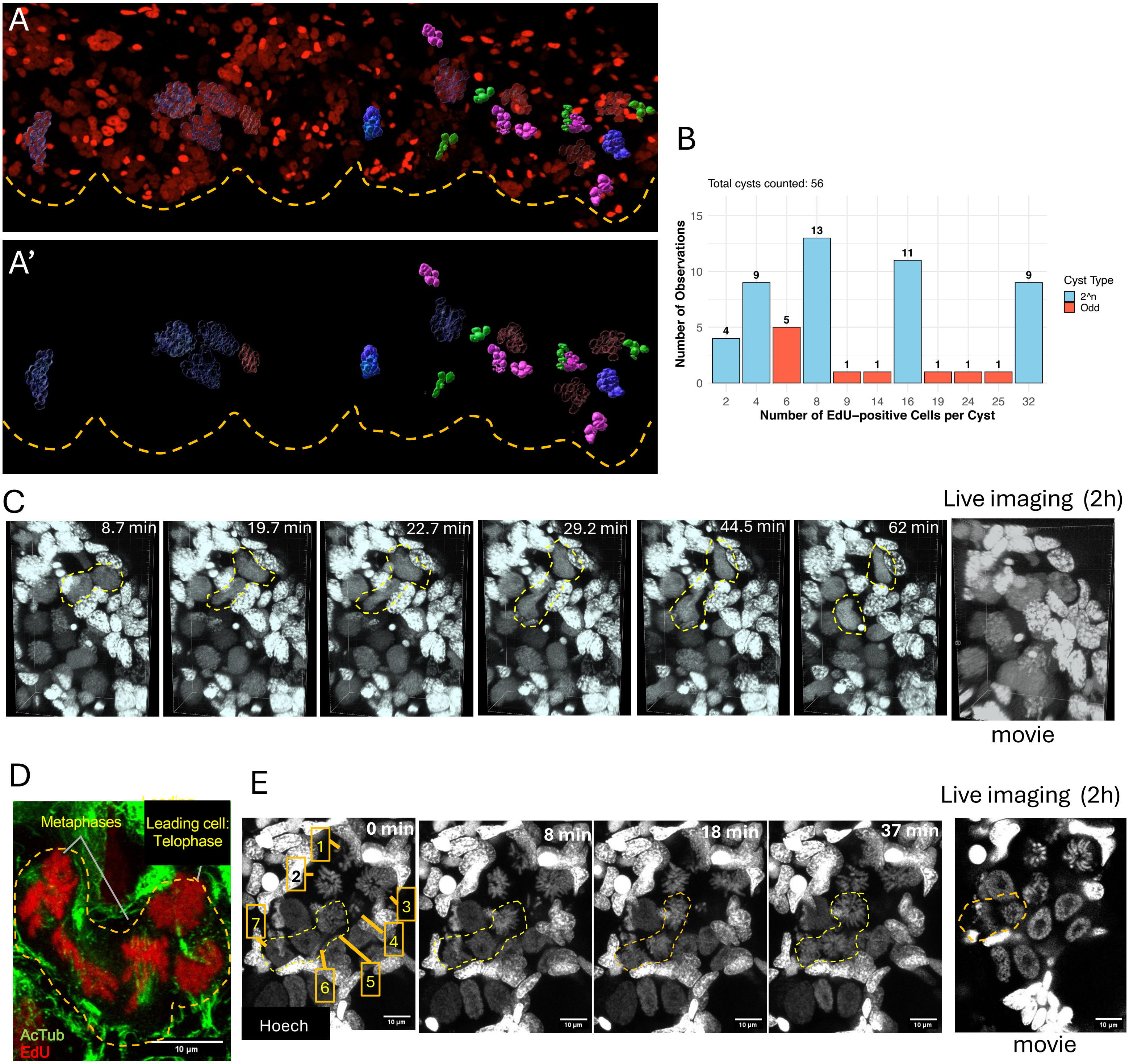

**Figure S4. Cyst dynamics and asynchrony.**

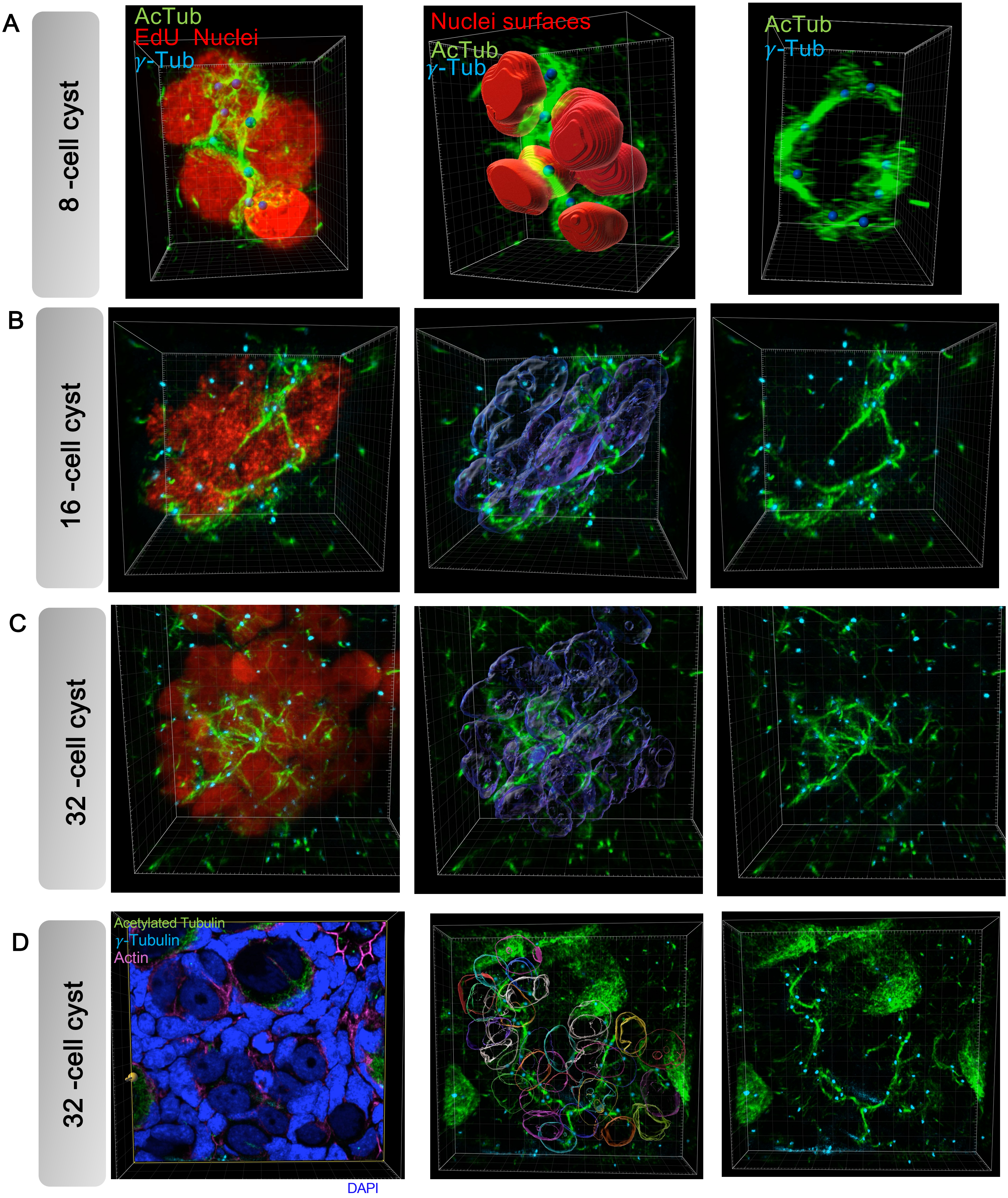

Figure S5. FLS in 8-, 16-, and 32-cell oogonia cysts visualized in Imaris as 3D (movies).

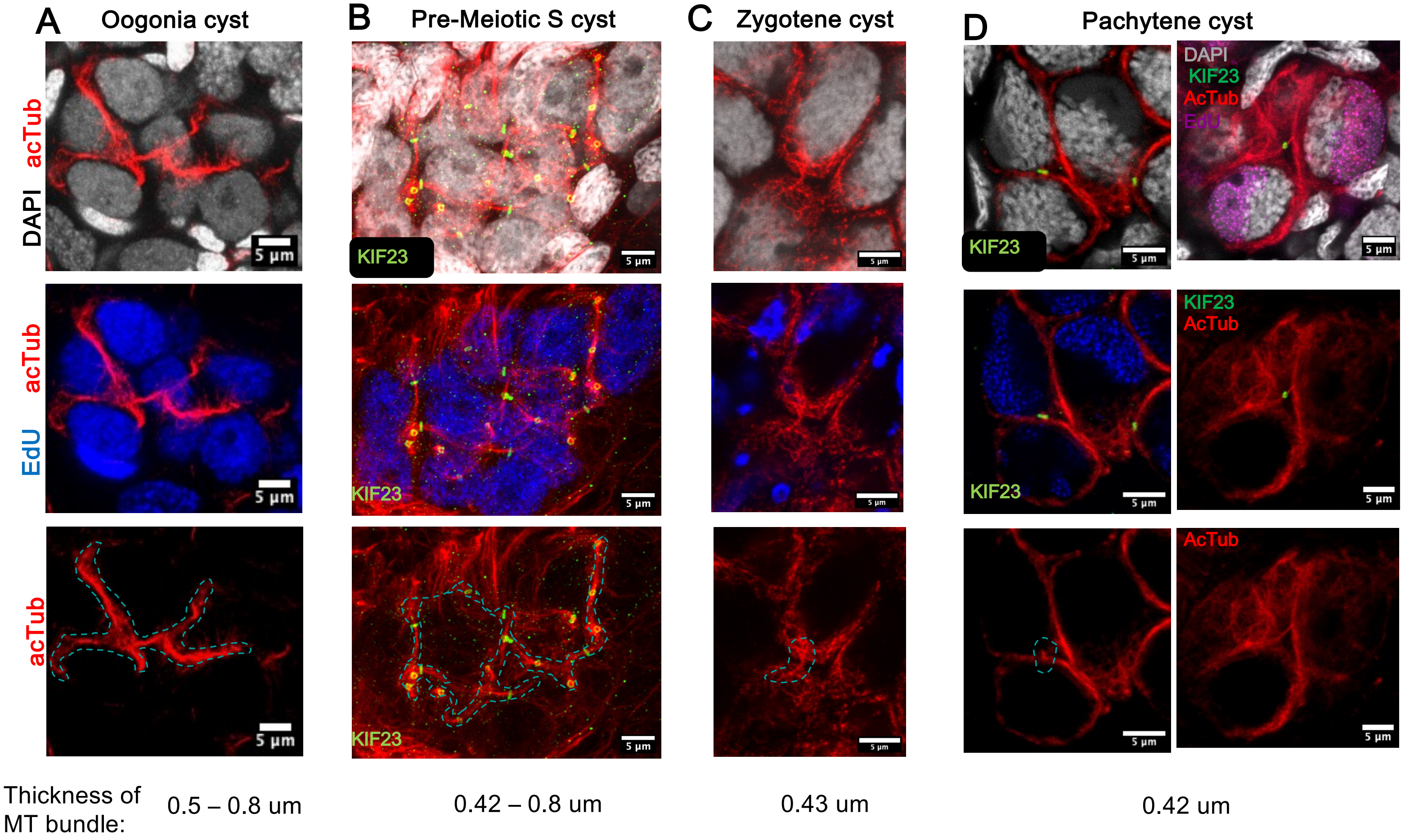

**Figure S6. FLS during germline cyst development from oogonia to pachytene.**

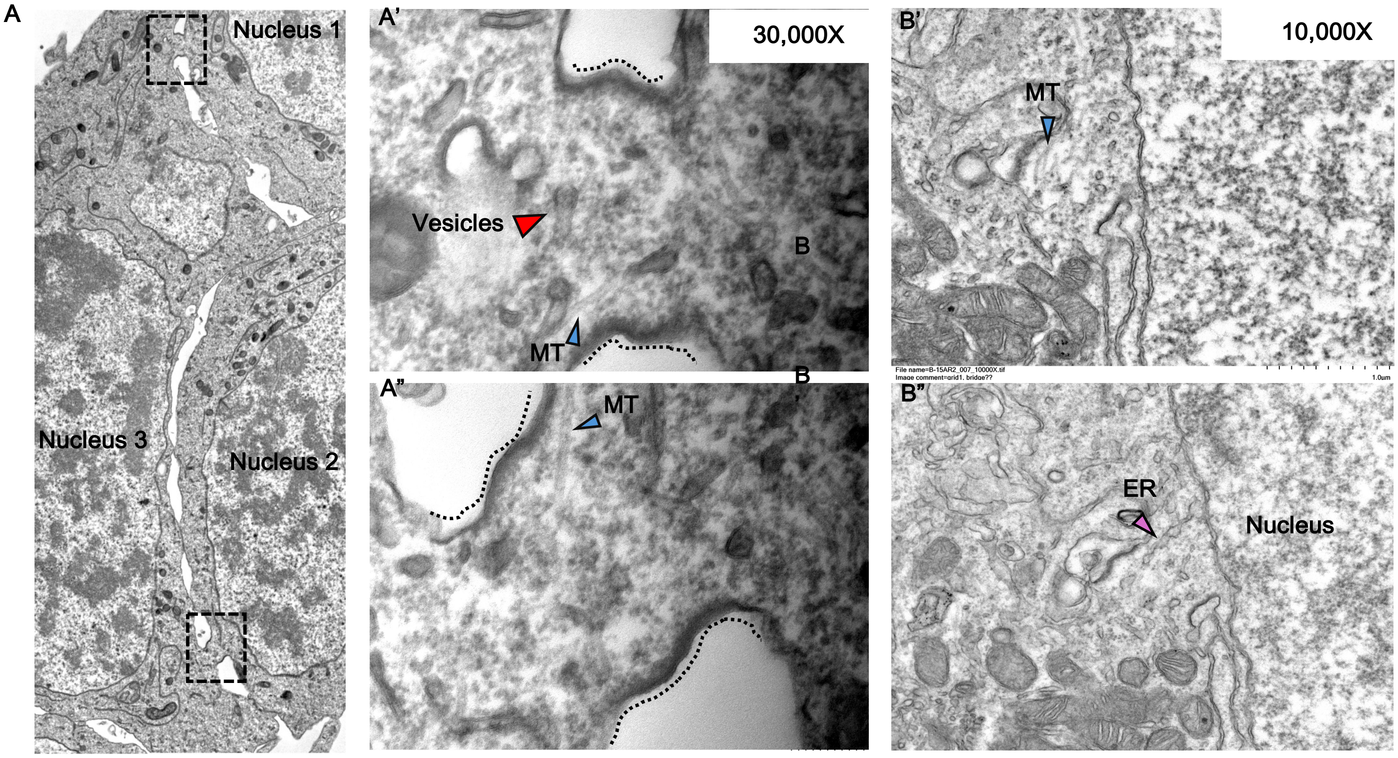

Figure S7. Ring canals with microtubules (MT), vesicles, and ER-like structures under electron microscopy (EM).

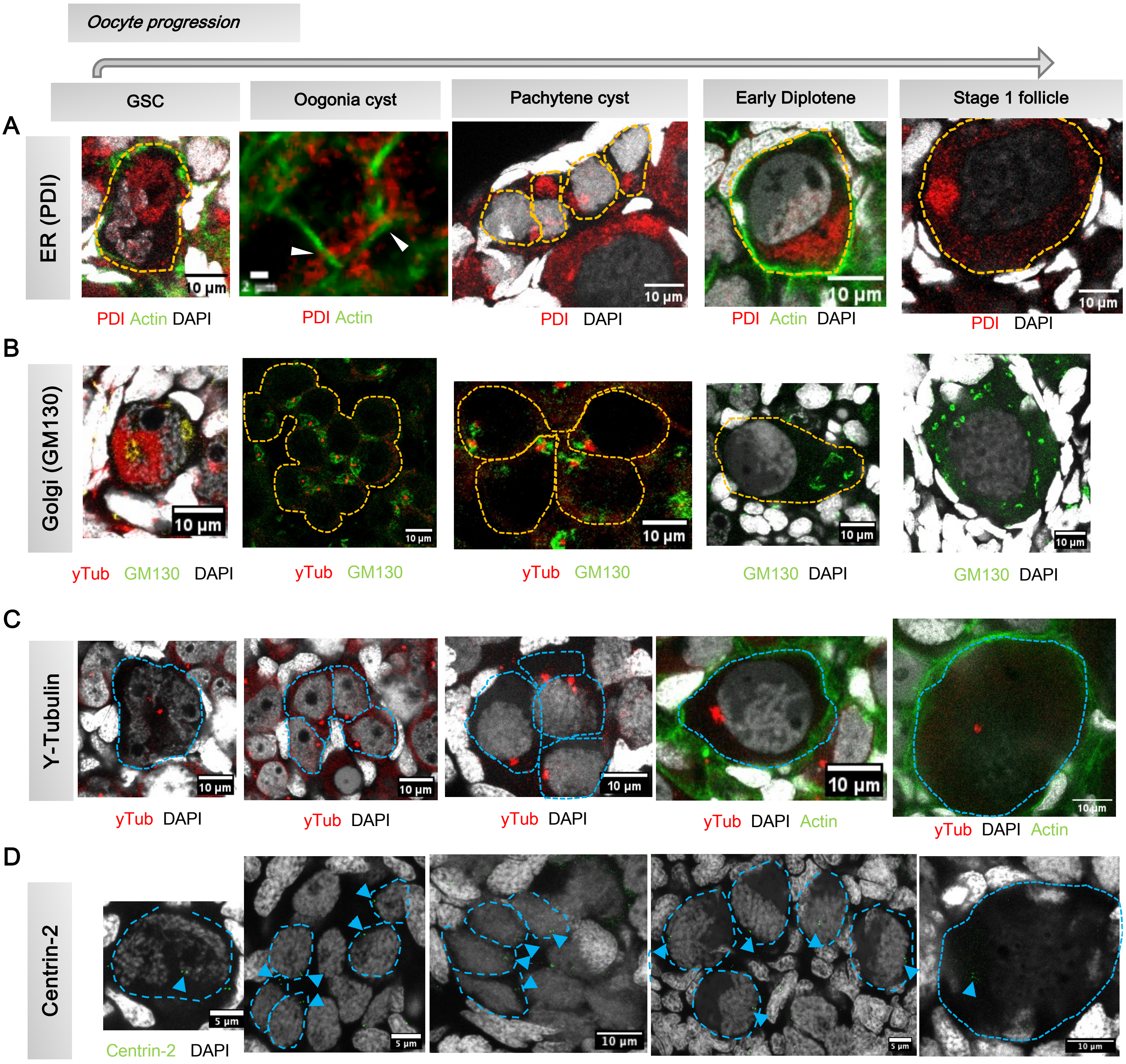

**Figure S8. Organelle dynamics during germline cyst development.**

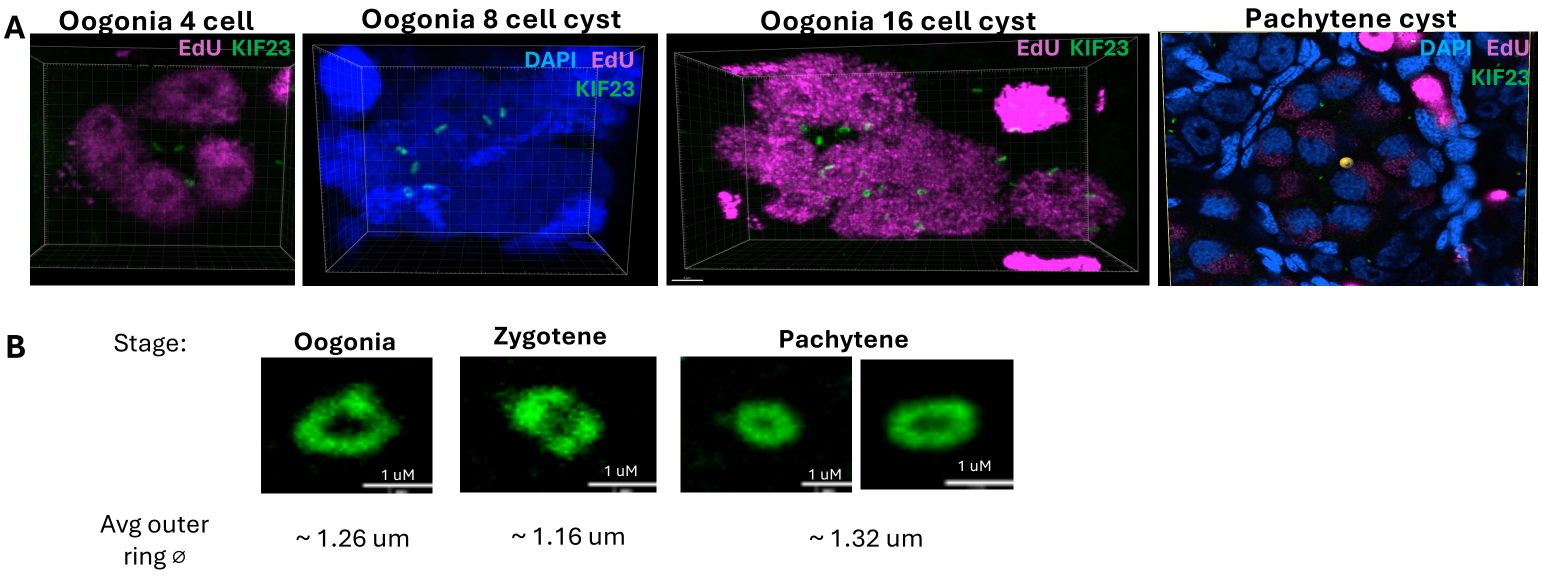

Figure S9. Ring canals during different cyst stages (movies).

Figure S10. Effect of Nocodazole treatment on cyst microtubules.

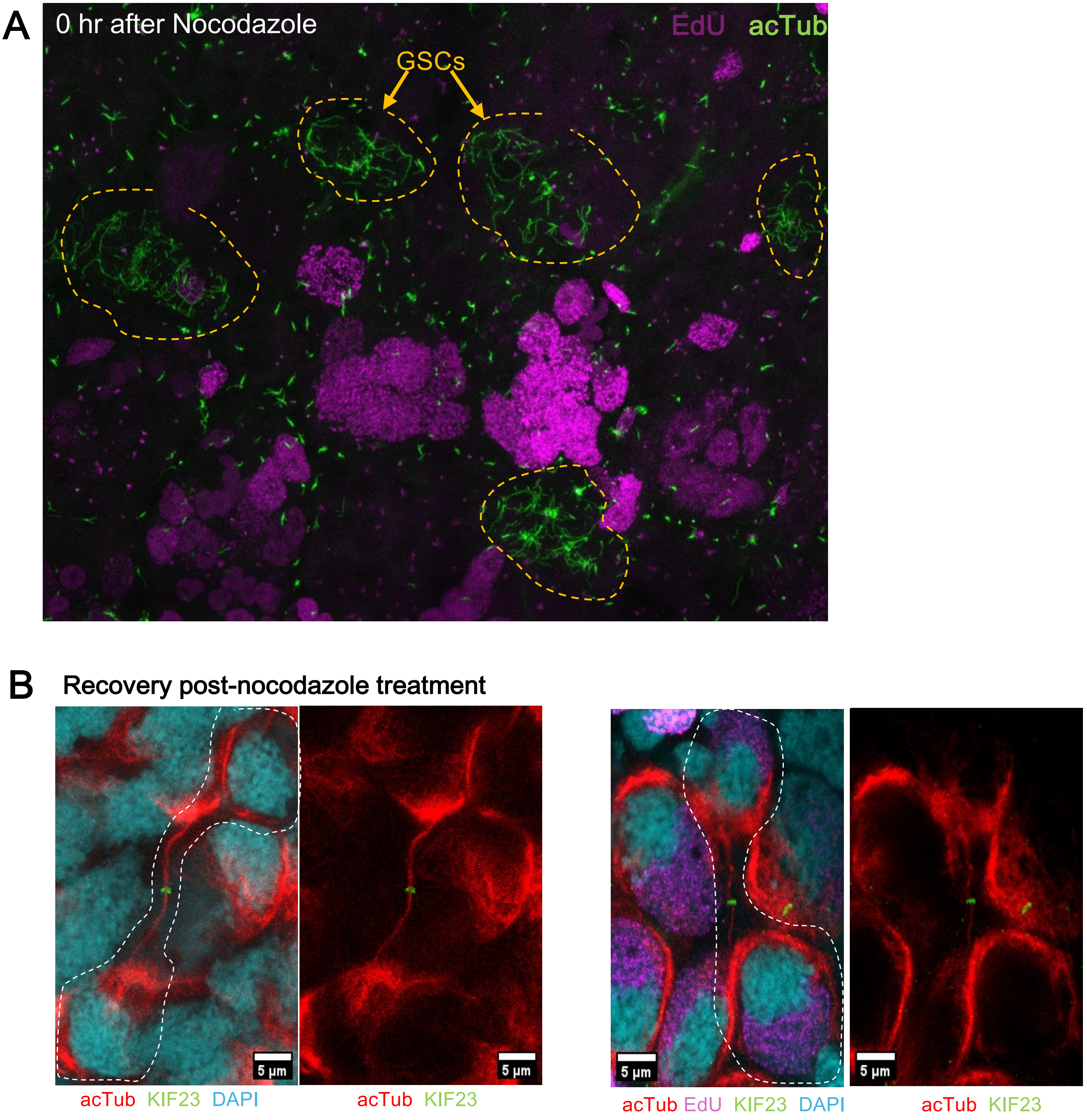

### Figure S11. Timeline of germline cyst development by EdU pulse.

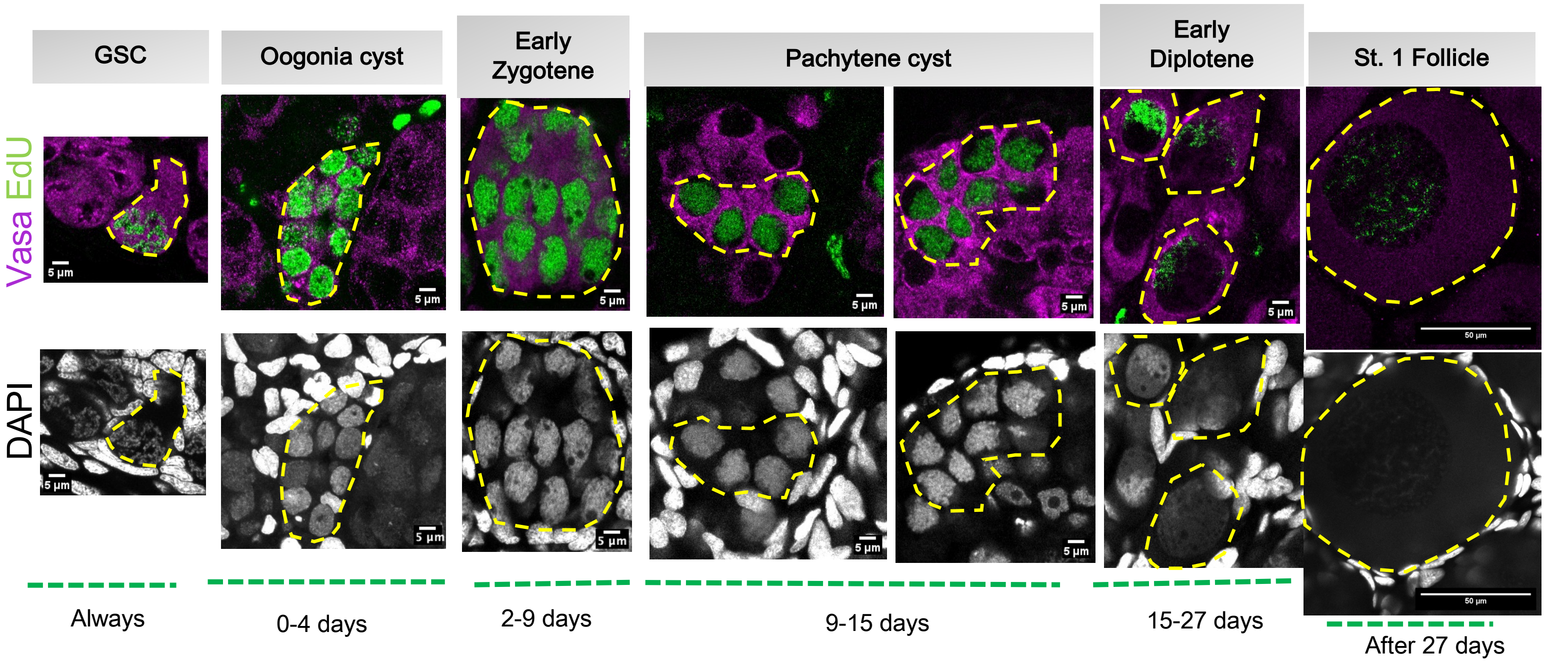

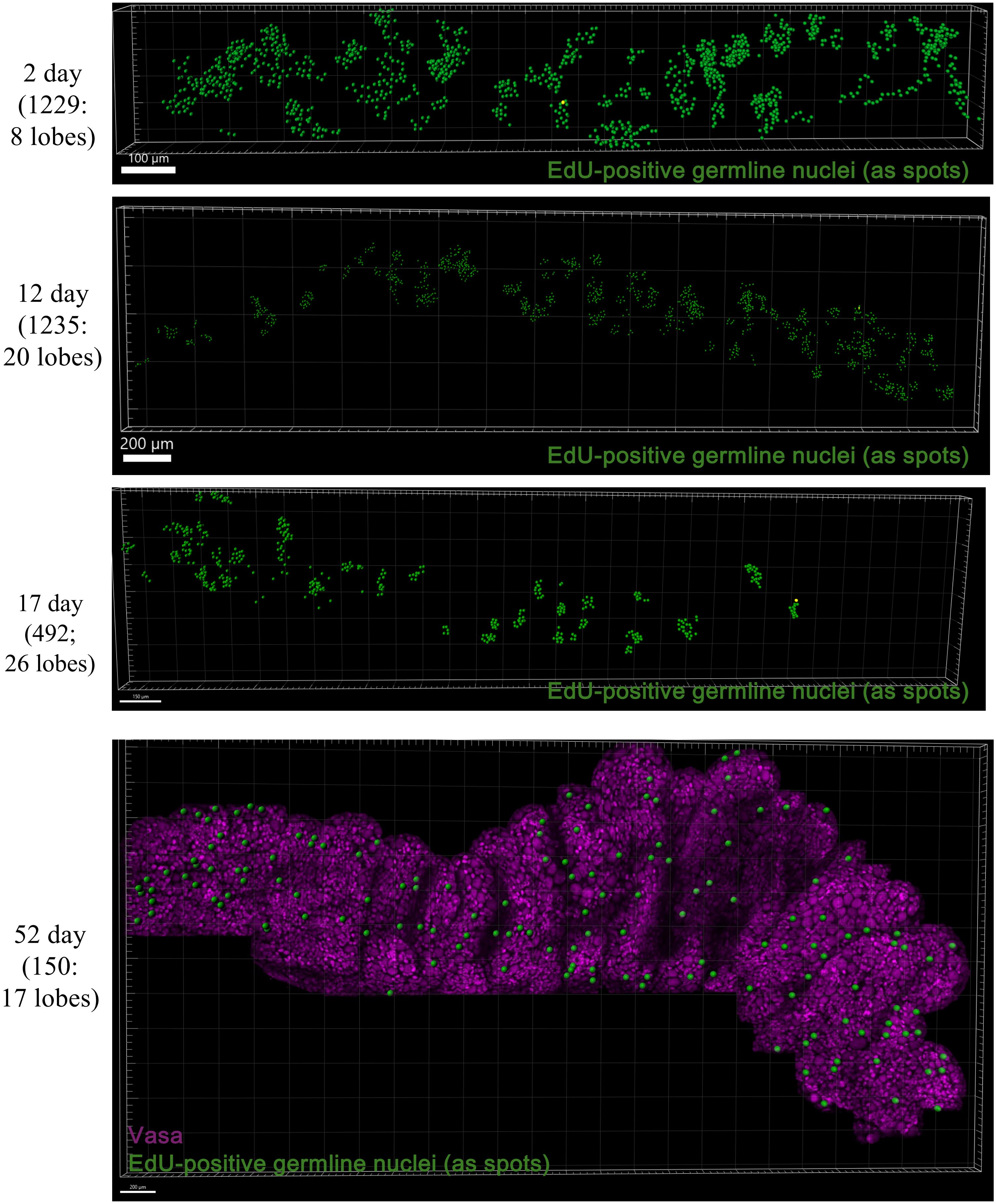

**Figure S12. Quantification of EdU-positive germline nuclei from EdU pulsed ovaries.**
